# Generative Design of New-to-nature Biosynthetic Assembly Lines with Genomic Language Modeling

**DOI:** 10.64898/2026.09.11.750945

**Authors:** Nathan Lanclos, Kyrellos E. Ibrahim, Andre Cornman, Marco Huang, Vikram Gill, Aalini Jain, Jonathan Abraham, Jennifer W. Gin, Yan Chen, Christopher J. Petzold, Justin Baerwald, Tanja Kortemme, Jay D. Keasling, Yunha Hwang

**Affiliations:** Joint BioEnergy Institute, Emeryville, CA, USA; Department of Bioengineering, University of California, Berkeley, CA, USA; Department of Bioengineering and Therapeutic Sciences, University of California, San Francisco, CA, USA; Biological Systems and Engineering Division, Lawrence Berkeley National Laboratory, Berkeley, CA, USA; Graduate Program in Chemistry and Chemical Biology, University of California, San Francisco, CA, USA; Tatta Bio, Cambridge, MA, USA; Computer Science Division, Department of Electrical Engineering and Computer Sciences, University of California, Berkeley, CA, USA; College of Computing, Data Science, and Society, University of California, Berkeley, CA, USA; Department of Molecular and Cell Biology, University of California, Berkeley, CA, USA; Department of Chemical & Biomolecular Engineering, University of California, Berkeley, CA, USA; Quantitative Biosciences Institute (QBI), University of California, San Francisco, CA, USA; California Institute for Quantitative Biosciences (QB3), University of California, Berkeley, CA, USA

**Author notes:** co-correspondence. These authors contributed equally (Nathan Lanclos, Kyrellos E. Ibrahim, Andre Cornman). Department of Biology, Department of Electrical Engineering and Computer Science, Schwarzman College of Computing, Massachusetts Institute of Technology, MA.

## Abstract

Reprogramming biosynthetic assembly lines can extend biosynthesis beyond the chemical space explored by nature. However, this remains difficult because assembly-line function depends on coordinated interactions across large multidomain enzymes. Here, we couple gLM2, a genomic language model trained on metagenomic sequences, with discrete diffusion and domain-level conditioning to enable generative design and optimization of biosynthetic gene clusters. We apply this approach to a chimeric type I polyketide synthase (PKS) engineered to produce δ-valerolactam, a molecule not naturally synthesized by PKSs. Through iterative redesign of two multi-domain regions in the context of the full PKS sequence, gLM2 progressively improved δ-valerolactam production, yielding variants with up to 9.4-fold higher titer than the starting enzyme. Together, these results demonstrate that evolutionary sequence information can be learned and applied to complex, multi-domain enzyme design problems, expanding biosynthetic assembly lines to produce molecules outside their natural biosynthetic repertoire.

## Main Text

Genomic language models (gLMs) trained on trillions of base pairs of diverse genomes learn evolutionary patterns governed by molecular and functional interactions across genomic elements^1–3^. Recent autoregressive gLMs^4^ have begun to leverage these learned dependencies to extend generative biology beyond individual proteins to multi-component systems^5^, most recently the de novo design of functional bacteriophages^6^. An open question is whether such models can be used not only to recapitulate natural function, but also to design large multi-gene systems that expand functions observed in nature. Here, we address this question by applying genomic language modeling to the programmable design of new-to-nature biosynthetic assembly lines.

Biosynthetic assembly lines are large multi-domain, multi-enzyme complexes encoded in biosynthetic gene clusters (BGCs) that span tens to hundreds of kilobases^7^ (Fig. 1a). They carry out assembly-line chemistry to produce structurally complex secondary metabolites, collectively referred to as natural products, which mediate diverse biological roles ranging from signaling to environmental stress response and chemical defense^8^. The same bioactivities that mediate these ecological functions have also made many natural products valuable as therapeutics, including antibiotics, antifungals, anticancer agents, and anti-inflammatory drugs. More than half of FDA-approved small-molecule drugs over the past four decades are natural products, their derivatives, or synthetic mimics thereof^9^. Beyond therapeutics, natural products are increasingly central to specialty chemicals such as dyes, flavors, and industrial additives^10^. More recently, biosynthetic assembly lines have been explored as a sustainable route to high-performance fuels^11^, materials^12^, and commodity chemicals^13^ that would otherwise rely on petrochemical feedstocks.

**Figure 1.**
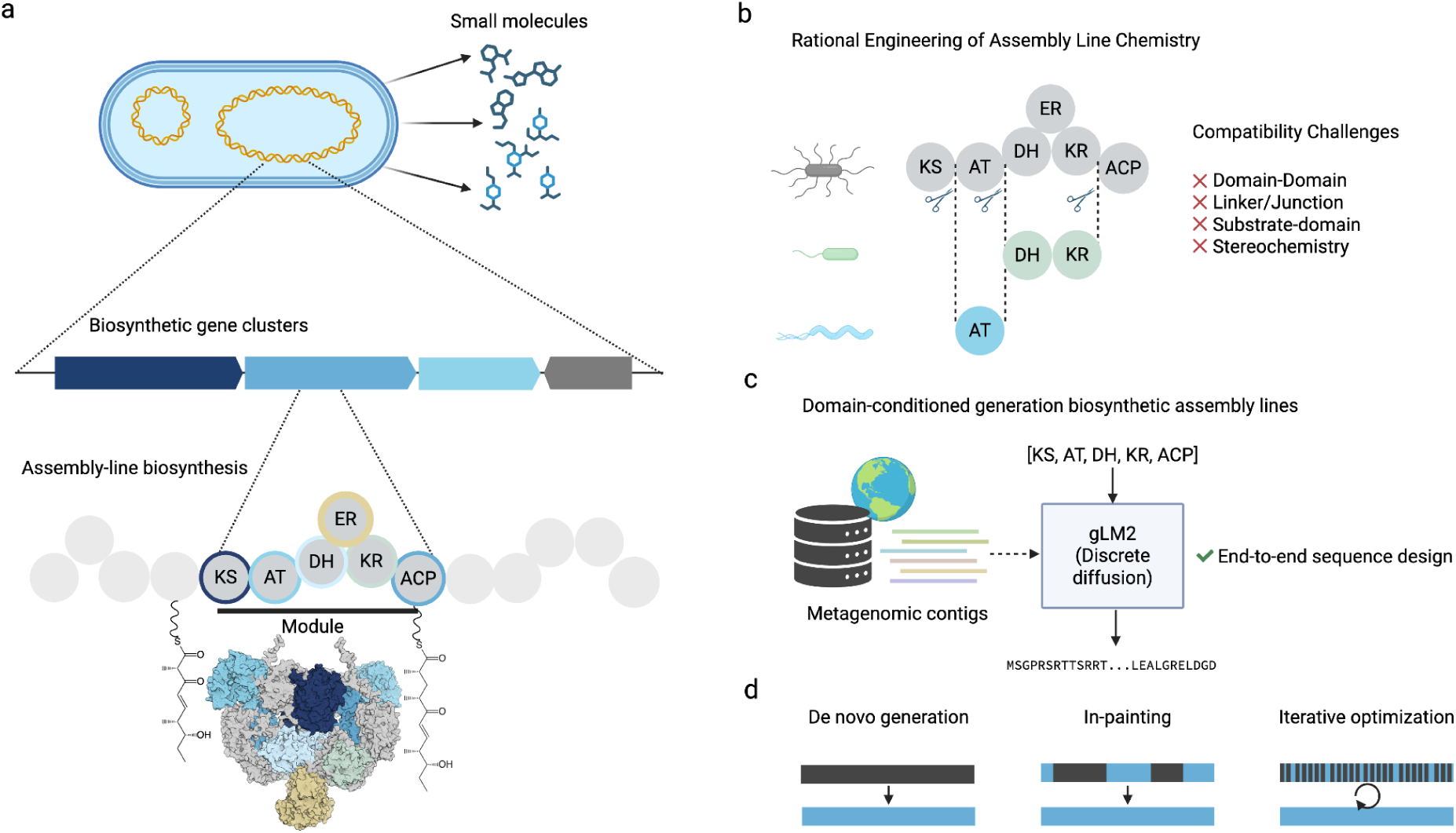
Genomic language modeling enables programmable design of biosynthetic assembly lines. **a.** Biosynthetic gene clusters encode multi-domain assembly lines that coordinate sequential enzymatic reactions to produce complex small molecules. Type I polyketide synthases (PKSs) are organized into catalytic domains whose order encodes biosynthetic logic. (KS, ketosynthase; AT, acyltransferase; DH, dehydratase; ER, enoylreductase; KR, ketoreductase; ACP, acyl carrier protein). **b.** Conventional PKS rational engineering recombines domains or modules from different natural assembly lines to alter product chemistry, but non-native combinations frequently exhibit poor functional compatibility. **c.** gLM2 is trained on large-scale genomic sequence data to learn evolutionary dependencies among protein sequences and their genomic context. Generation is conditioned on a specified architecture of catalytic domains, enabling controllable generation of protein sequences consistent with the desired biosynthetic domain order. **d.** Coupling gLM2 to discrete diffusion enables sequence generation under different levels of conditioning, including de novo generation, template optimization, and sequence in-painting while preserving specified sequence features. Figure created in BioRender.

In nature, biosynthetic assembly lines diversify primarily through recombination of modular domain architectures^14,15^. This natural mode of diversification has motivated decades of engineering effort to programmably synthesize desired molecules by combinatorially mixing and matching modules and domains across assembly lines (Fig. 1b). The plug-and-play vision of building a biological assembly line that synthesizes any target molecule on demand is particularly compelling because total biosynthesis promises stereospecific, enantiopure, and efficient routes that are difficult to achieve with chemical synthesis^16^. In practice, however, rational engineering by domain or module swapping remains slow, low-yielding, with outcomes that are difficult to predict because the biophysical rules governing module compatibility and interdomain communication are still poorly understood. Recent works have shown that evolution-guided identification of fusion sites^17,18^, namely junctions that mirror natural recombination boundaries, can improve chimeric assembly line construction. Yet, because multi-step catalysis depends on coordinated substrate channeling and interdomain contacts^19,20^, even evolutionarily informed chimeras often remain functionally compromised.

We hypothesized that a genomic language model that has internalized the evolutionary statistics of natural BGCs would capture the implicit grammar of compatible domain and module pairings, and could therefore design more functional chimeric assembly lines. We build on gLM2^2^, a mixed-modality genomic language model trained on a large metagenomic corpus to jointly model protein-coding and intergenic sequences (Fig. 1c). Because gLM2 was built as a masked encoder, it does not natively support generative sampling. In this work, we make gLM2 generative by coupling it to a discrete diffusion framework that supports in-painting, template optimization, and de novo generation (Fig. 1d). The first two regimes are particularly suited to assembly line engineering, where defined catalytic domains must be preserved while flanking and linker sequences are redesigned. We further make this generative model controllable by conditioning on biosynthetic domains. We focus our demonstration on Type I polyketide synthases (PKSs) (Fig. 1ab), a versatile class of assembly lines in which biosynthetic logic is colinearly encoded in the domain order of the sequence^21^. PKS modules are minimally composed of a ketosynthase (KS), an acyltransferase (AT), and an acyl carrier protein (ACP) that coordinate for chain elongation, and can include additional processing domains: a ketoreductase (KR), dehydratase (DH), and enoylreductase (ER) that sequentially reduce the β-carbon. Specifically, we use gLM2 to redesign large multi-domain segments in a chimeric PKS engineered to produce a C5-lactam used in nylon polymerization^22,23^, a product not naturally synthesized by PKSs (Fig. 2ab)^24^, and show that gLM2 designs iteratively improve product titers up to 5.7-fold and 1.67-fold across sequential design rounds.

## 2. Results

### 2.1. Coupling gLM2 with discrete diffusion enables programmable design of biosynthetic gene clusters

gLM2 is a mixed-modal genomic language model trained on the OpenMetaGenomic (OMG) database, comprising 3.1 trillion base pairs of metagenomic sequences^2^. gLM2 takes as input genomic sequences that are pretokenized via gene-calling such that protein-coding genes are represented as amino acid sequences while intergenic regions are represented as nucleic acid sequences. Initially, gLM2 was trained using a masked language modeling objective with a fixed 30% masking rate. We previously demonstrated that gLM2 internalizes co-evolutionary signals across protein-protein interactions, a property that can be leveraged to predict protein-protein interaction interfaces at residue-level resolution^25^.

To harness these co-evolutionary signals for the design of large multi-protein complexes, we adapted gLM2 for sequence generation by finetuning on 6.2 million microbial contigs (comprising 400 million proteins) from the OpenGenome (OG) database^2^ using a masked discrete diffusion modeling framework^26^. Intergenic regions were removed to focus the model’s generative capacity exclusively on protein sequences (Fig. S1a, Fig. S2). Unlike the base gLM2 model, which used a fixed masking rate for representation learning, the generative diffusion approach required the model to iteratively denoise sequences across all possible noise levels. Accordingly, the model was trained to minimize cross-entropy loss under a dynamic masking scheme where the masking rate for each coding sequence was independently sampled between 0% and 100%. This flexible framework enables both unconditional sequence generation (100% masking) and context-conditioned design, allowing partial sequence information, such as neighboring proteins or fixed functional domains, to guide the generative process.

We further finetuned gLM2 on the BiG-FAM database^27^ using the same discrete diffusion framework. This dataset consists of 1,225,071 biosynthetic gene clusters, including 211,243 PKSs, 337,201 non-ribosomal peptide synthases (NRPSs), and 79,506 PKS-NRPS hybrid clusters. (Fig. S1b, Fig. S2). To accommodate these large multi-protein systems, we increased the model context window to 16,384 amino acids (Fig. S3). Additionally, we implemented domain-level conditioning by utilizing biosynthetic domains curated using antiSMASH^28^. Specifically, the model vocabulary was expanded to include 1,999 unique domain families as discrete special tokens (e.g. <KETOACYL-synt>, <KR>) which were prepended to the corresponding protein coding sequence and excluded from the masking process, allowing them to serve as fixed conditioning prompts. This architecture enables the generation of biosynthetic gene clusters prompted by a specified list of domains during inference, effectively providing programmable control over biosynthetic order.

### 2.2. Optimizing a Chimeric PKS for Valerolactam Production

To investigate the ability of the fine-tuned gLM2 model to design a biosynthetic assembly line, we set out to improve an existing chimeric PKS sequence via template optimization. A rationally engineered chimeric PKS that produces δ-valerolactam (VL)^24^ was selected as a model system with established experimental protocols, enabling straightforward evaluation of variants generated by gLM2. The template PKS, hereafter referred to as VL-Start, joins the loading module and catalytic KS domain of the Fluviricin PKS module 1^29^, the second extension module of Cremimycin PKS^30^ (starting at the KS adapter domain at C-terminus of the KS^31–33)^, and the native Flu TE (Fig. 2a, Fig. S4a). VL-Start has two non-native junctions: at the C-terminus of the KS at the structured adapter domain^33^ and between the ACP of the extension module and the Flu TE (Fig. S4a). Notably, VL-Start was the highest VL-producing PKS of a library of 140 module exchanges with fusion sites across the entire Flu KS domain and KS-AT linker, consistent with evolutionarily informed module boundaries and related engineering successes (Fig. S4b)^34–38^.

**Figure 2.**
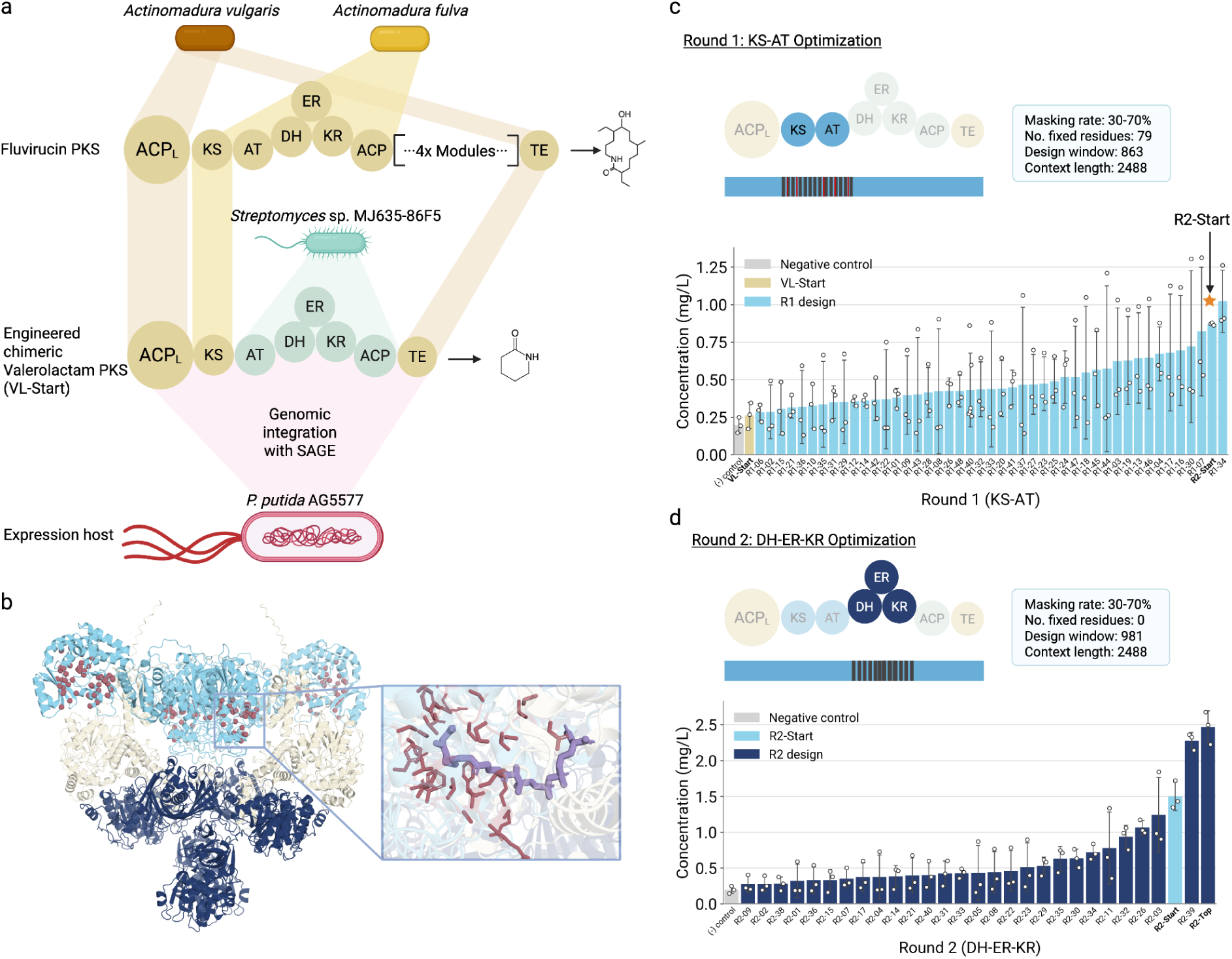
Sequential context-conditioned redesign of a chimeric PKS increases valerolactam production. **a.** Schematic of the chimeric PKS used as the starting sequence (VL-Start) and genomic integration of engineered PKSs into the *Pseudomonas putida* production strain using Serine Recombinase-assisted Genome Engineering (SAGE). The engineered assembly line produces δ-valerolactam. **b.** AlphaFold3-predicted structure of VL-Start used to define redesign boundaries. The inset shows predicted ligand-interacting residues used as fixed positions during Round 1 generation. **c.** Round 1 redesign of the KS-AT region. gLM2 sequences were generated at 30-70% masking while preserving 79 fixed residues within an 863-residue design window. Valerolactam titers are shown for experimentally tested variants ranked by production. The starting sequence for the Round 2 (DH-ER-KR) optimization was selected from one of the top performers (R2-Start). **d.** Round 2 redesign of the DH-ER-KR domains using the Round 1 top-performing sequence (R2-Start) as the template (Supplementary Note 1). No residues were fixed within the 981-residue design window. Individual points are technical replicates (n = 3), variants are ranked by valerolactam production (mean ± SD). Figure created in BioRender.

Biosynthesis of VL proceeds by loading a preprocessed aminoacyl starter unit onto the KS domain, followed by a 2-carbon extension via condensation with malonyl-CoA (M-CoA) preloaded onto the downstream ACP by the AT domain. The β-carbon ketone of the ACP-bound extended polyketide is then fully reduced by the β-carbon processing domains (DH, ER, KR). Following deprotection by an accessory enzyme, the TE of the final Fluvirucin PKS module hydrolyzes and cyclizes the extended and processed polyketide chain into δ-valerolactam (Fig. S5)^24^. Production of this and any polyketide product requires careful coordination of the downstream ACP with each catalytic domain in the assembly line, and thus, context-aware design is required to reconcile structural and catalytic compatibility across the assembly line and its catalytic cycle^39–45^. Because the penultimate synthesis step deprotects a highly reactive primary amine, we reasoned that the ACP::TE interaction is unlikely to be rate-limiting for production and prioritized the KS-AT junction in the first round of designs.

We developed our design strategy in two phases aiming to 1) improve the chimeric KS-AT didomain of the PKS and 2) optimize the downstream processing domains for compatibility with the redesigned didomain. While structurally proximal, the KS and AT catalyze distinct steps that are coupled through the downstream ACP^41,46,47^. We reasoned that sequence optimization will primarily improve non-native contacts at the chimeric interface and the overall stability of the KS-AT didomain, while maintaining coordination with the ACP for the transacylation and condensation reactions. In fully reducing PKSs, successful product formation requires correct processing to modify the oxidation state of the β-carbon (in order: KR reduces β-ketone to β-hydroxyl, DH dehydrates to alkene, and ER reduces to alkane)^45^. If the binding or catalytic capacity of any upstream domain is disrupted, the functionality of the downstream domain(s) are insufficient to rescue production of the correctly reduced target molecule. Context-aware sequence design in theory should be able to coordinate residue assignment across multiple domains, coupling the enzymatic constraints with the protein-protein interaction requirements for successful catalysis. Separating our design goals into two successive rounds derisked the scope of the design campaign, avoided the cost of long (7.5kb) and high-GC (∼60%) DNA synthesis, and enabled us to evaluate the performance gains associated with each stage of redesign.

### 2.3. Context-conditioned constrained design of the KS-AT didomain improves non-natural lactam synthesis

We hypothesized that the highest improvement in yield would come from first optimizing the local structural compatibility at the chimeric KS-AT didomain that coordinates extender-unit selection, substrate loading, and decarboxylative chain elongation in conjunction. Using a structure-informed approach, we defined the redesign boundaries by selecting the N-terminal KS residue and C-terminal AT residue that belong to the structured, globular catalytic domains determined by an AlphaFold3 (AF3) predicted structure of VL-Start (Fig. 2b). KS domains commonly act as substrate gatekeepers that discriminate incoming polyketide intermediates^48^. To preserve catalysis and substrate binding to account for KS gatekeeping or AT selection of M-CoA, the active site residues were fixed during gLM2 sequence generation (Fig. 2bc). Specifically, active site residues were determined by predictive co-folding of each individual domain with its respective ligand (KS: β-ketophosphopantetheine, AT: malonyl-CoA) with Boltz-2^49^. We assessed the reconstruction accuracy of Boltz-2 on PKSs by refolding 24 solved structures of partial PKS modules containing biologically relevant ligands (Table S1, Fig. S6). KS and AT ligand placement success (<2 Å heavy-atom RMSD) was 38% and 85% (Fig. S6a), respectively, and always correctly identified the pocket.

To generate a diverse cohort of candidates, we sampled sequences using the fine-tuned gLM2 by randomly masking residues within the design window with masking rates 0.3, 0.5, and 0.7 and reconstructing them via discrete diffusion. The resulting sequences were evaluated by gLM2 pseudo-likelihood scores and filtered by Shannon entropy to remove low-complexity outputs. From this filtered distribution, 45 KS-AT designs were selected for experimental validation by sampling evenly across the pseudo-likelihood score distribution and across the different masking rates.

Codon-optimized DNA sequences encoding the design region were synthesized and assembled into plasmids encoding the full length PKS sequences. PKS genes were chromosomally integrated into the previously described Serine-Recombinase Assisted Genome Engineering (SAGE) compatible *Pseudomonas putida* strain LM3-FlvJ (Fig. S7)^24,50^. The KS-AT design library was evaluated for VL production by cultivation and liquid chromatography-mass spectrometry (LCMS) analysis for quantitative product titers (Fig. S8, Methods).

In Round 1 (KS-AT) optimization, all 45 designs matched or improved the VL production of VL-Start, with 84% (38/45) exceeding one standard deviation above the VL-Start titer. The highest producing variant (R2-Start) produced approximately 5.7-fold more VL than VL-Start (Fig. 2c) across multiple independent cultivations (Fig. S9-S10, Supplementary Note 1). R2-Start was subsequently selected as the new template for the redesign of the remaining catalytic domains (DH-ER-KR).

### 2.4. Context-conditioned unconstrained design of the β-carbon processing domains further enhances non-natural lactam synthesis

Having established in Round 1 that constrained, structure-guided generation could improve VL production, we next asked whether gLM2 could redesign a catalytic region without predefined residue-level constraints. The Round 1 strategy preserved residues predicted by Boltz-2 to contact the KS and AT substrates, reducing the risk of disrupting known catalytic functions but also introducing assumptions about which positions must remain unchanged. Such constraints may limit the generality of a generative design framework, particularly when high-confidence ligand-bound structures are unavailable or experimentally defined catalytic determinants are not well-defined. Even when the chemical structure of the substrate is known, as in the VL pathway, the sequence features required for its recognition and processing cannot necessarily be specified from a predicted binding pocket alone. More broadly, enzyme substrate specificity can depend on residues distributed beyond the immediate catalytic site^51^. This limitation is particularly relevant to less characterized or engineered biosynthetic assembly lines, for which substrates, products, and catalytic determinants are frequently not experimentally resolved or native to the wildtype sequence. Experimentally determined reduction domain data are sparse, with only a handful of relevant ligand-containing structures in the PDB (13 total, 8 co-factor only) (Table S1). Due to this, DH and ER domains are especially not well represented, and the docking evaluation reflected this with 18.2% success (<2 Å heavy-atom RMSD) rates (Fig. S6a). Both unconstrained and constrained docking (see methods) on solved multi-domain structures had a 0% success rate with significant decreases in contact recovery (Fig. S6c, S6f). We therefore allowed end-to-end generation of the complete set of β-carbon processing domains to test whether gLM2 could autonomously preserve sequence features required for catalysis.

We selected the β-carbon processing domains DH-ER-KR as a stringent test of unconstrained, context-aware design because it couples three sequential catalytic transformations on the same ACP-bound intermediate. VL synthesis requires the three domains to accommodate successive chemical states of the non-native intermediate while maintaining compatibility with the ACP and with one another. Previous reductive-loop engineering has shown that successful transfer depends strongly on compatibility between the loop and the chemical structure of the substrate it encounters^52^, while structural studies of PKS reducing regions indicate that substrate recognition involves features extending beyond the canonical catalytic residues^45,53^. We hypothesized that gLM2 could circumvent the lack of reliable structural priors by learning which sequence features must be preserved for reductive catalysis while simultaneously exploring sequence space to improve overall production of the non-native lactam intermediate. Such capability would be necessary for applying the framework to less-characterized biosynthetic assembly lines, where the structural information needed to define residue-level constraints is often unavailable.

Using R2-Start as the template, we generated 1,500 sequences for the DH-ER-KR region using masking rates of 0.3, 0.5, and 0.7 without fixing any residues within the design window. We found that gLM2 conserved identified catalytic residues and preferentially retained active site residues in unconstrained sampling (Fig. S11, Supplementary Note 2)^54–58^. Twenty-eight generated designs were selected for synthesis, cloned, and tested following Round 1 methods with the principal experimental difference being the use of shifted integration loci (Supplementary Note 1). The highest producing designs roughly doubled VL production over R2-Start, and produced up to 9.4-fold more VL than the rationally designed VL-Start PKS sequence (Fig. 2d). Proteomic analysis of the fermented strains indicate PKS expression is one driver of VL titer, but is insufficient to fully explain high or low producing designs (Fig. S12).

### 2.5. Interpretable gLM2 features capture local catalytic sites and 3-dimensional inter-domain proximity

To examine the sequence dependencies learned by gLM2, we can calculate categorical Jacobians^59^, which measure how changes at each residue affect the model’s predictions at every other position. We first asked whether these dependencies reflect physical, functionally important inter-domain connections, computing it directly on the sequences of three experimentally solved homologous PKS regions and measuring the coupled pairs in the deposited structures. The strongest long-range couplings (|J| > 4) are close inter-domain distances (<12 Å) in the DEBS KS-AT didomain (PDB 2QO3)^32^, the juvenimicin reducing region (PDB 8G7W)^45^, and the intact Lsd14 module (PDB 7S6B)^60^ (Fig. S13, Supplementary Note 3). gLM2 reliably identified interfaces of documented importance: the KS-AT linker/post-AT-linker interface, whose boundary also governs functional AT-domain exchange^61^, and the DH-ER-linker (KR_S_)/KR interface^45^. Whereas Multiple Sequence Alignment (MSA)-based analyses can identify coevolving positions with sufficiently deep and diverse sequence sets, gLM2 can generatively sample sequence variants of a single sequence input that preserve learned constraints across the assembly line.

We then applied the same analysis to the starting PKS sequence (VL-Start). The resulting map contained strong local dependencies within individual domains, as well as discrete long-range couplings between residues separated by hundreds of positions in the primary sequence (Fig. 3a). Mapping these couplings onto the AlphaFold3-predicted structure showed that many corresponded to predicted close distances between domains and linkers. Notable examples included interactions between the KS-AT and post-AT linkers and between the DH-ER linker (KRs subdomain) and the KR domain (Fig. 3b). Thus, the model’s internal sequence dependencies recover features of the higher-order organization of the assembly line, including interfaces that connect nonadjacent regions of the sequence.

**Figure 3.**
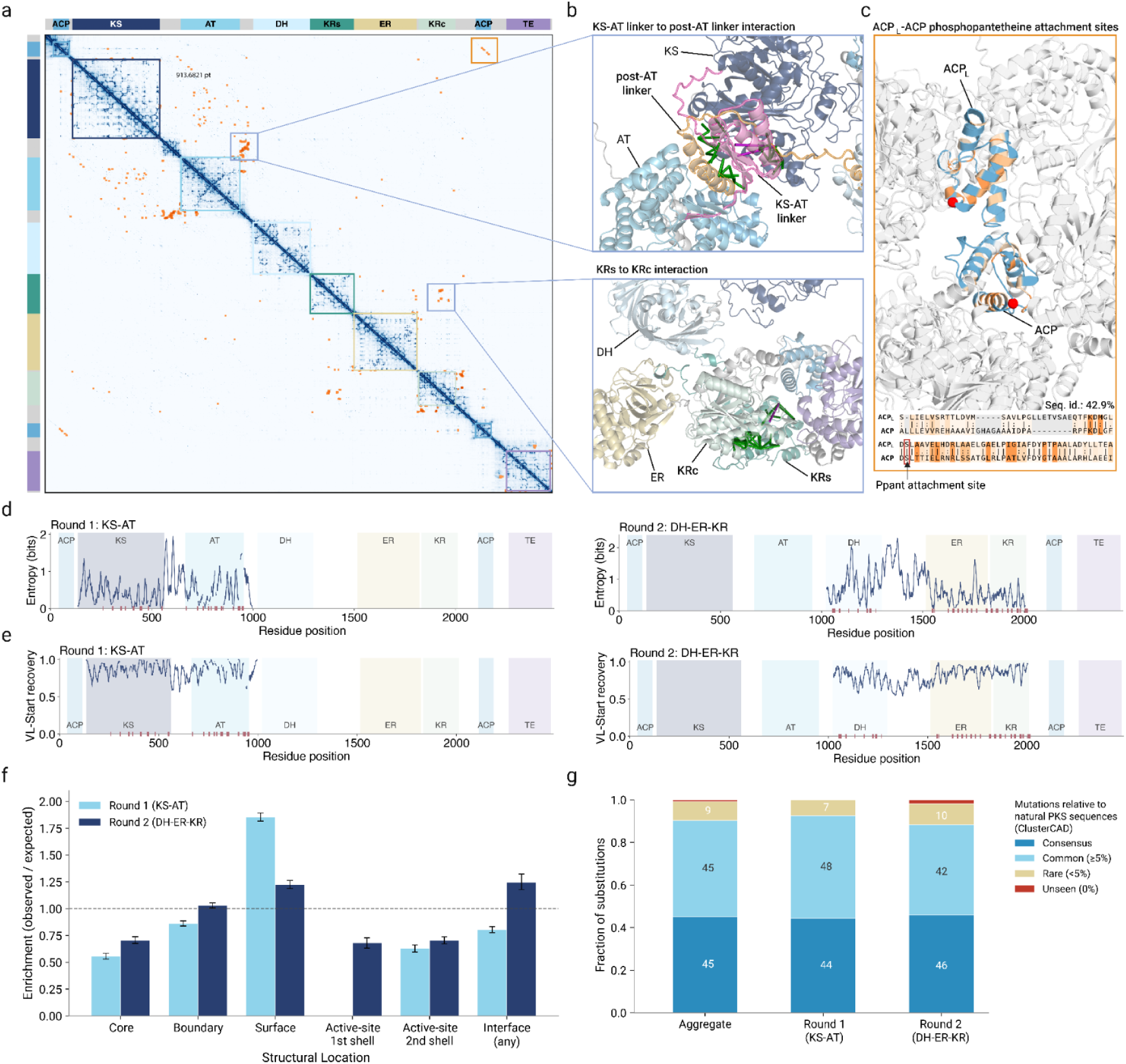
gLM2 captures structural, catalytic, and long-range dependencies within the PKS assembly line. **a.** Categorical Jacobian of the full PKS sequence showing position-wise dependencies learned by gLM2. Domain boundaries are indicated along both axes; top 0.01% long-range non-intra-domain couplings are highlighted in orange. **b.** Top 30 long-range predicted couplings in selected inter-domain regions mapped onto the AF3-predicted PKS structure. Highlighted interactions include couplings between the KS-AT linker (pink) and post-AT linker (orange) and between the KRs (structural subdomain between DH and ER) and KRc (catalytic subdomain). Green rods between true positive predicted Cβ-Cβ distances <12Å in predicted structures, and fuchsia rods are false positive (>12Å) in the AF3 model; the same interfaces are recovered in experimentally solved homologous PKS structures (Fig. S13). **c.** Long-range coupling between the loading ACP (ACP_L_) and downstream ACP regions surrounding their phosphopantetheine (Ppant) attachment sites (red sphere). Corresponding regions are shown on the predicted structures and sequence alignment. **d.** Position-wise sequence entropy across generated Round 1 KS-AT (left) and Round 2 DH-ER-KR (right) designs. Domain boundaries are shaded; ticks indicate fixed ligand-interacting residues in Round 1 and predicted ligand-interacting residues in Round 2 (see Methods). **e.** Native-sequence recovery across the two design windows, defined as the fraction of generated sequences retaining the corresponding residue from the starting template at each position. **f.** Enrichment (observed/expected) of gLM2 designed substitutions by location in the AF3-predicted structure. Burial categories defined by Rosetta Cβ-neighbor count. Interfaces defined as interdomain or interchain contacts. Error bars represent 95% CI across all tested sequences. **g.** Designed substitutions classified against natural PKS sequences (ClusterCAD) at each aligned position: consensus (most frequent residue), common (≥5%), rare (<5%), unseen (0%). Labels are percentages.

Not all long-range dependencies corresponded to close physical distances. gLM2 also coupled residues surrounding the phosphopantetheine attachment sites of the N-terminal loading ACP (ACP_L_) and the downstream ACP domain (Fig. 3c). These regions are spatially separated in the predicted structure, but are homologous (42.9% sequence identity), and the coupled residues fall within conserved structural and functional motifs that mediate phosphopentatheine attachment and substrate transfer along the assembly line. Their coupling therefore likely reflects a functional constraint local to the phosphopentatheine-attachment site within ACPs rather than structural proximity, consistent with the notion that gLM2 captures dependencies arising from conserved and coordinated catalytic roles in addition to structural proximity.

We next quantified position-wise sequence entropy across the generated designs to determine where the model permitted sequence diversification. Entropy varied substantially across each design window rather than being uniformly distributed (Fig. 3d). In the Round 1 (KS-AT) designs, the greatest diversity was generally observed in linker and peripheral regions. In the Round 2 (DH-ER-KR) designs, for which active site residues were not fixed during generation, predicted active site positions nevertheless showed markedly lower sequence entropy and were recovered more frequently than surrounding positions, corresponding to a 38-fold lower median substitution rate than the remainder of the design region (Fig. S11a). Consistent with this result, recovery of the starting-sequence residue varied at single-residue resolution and did not reveal extended sequence blocks that were consecutively identical to the template (Fig. 3e). Instead, annotated catalytic residues were unchanged in every designed and tested sequence (Fig. S11, Supplementary Note 2). This indicates that the model preferentially conserved catalytic constraints even when they were not supplied explicitly during sampling.

We next examined where the generated substitutions were located in the predicted structure and how they related to natural sequence variation. In both design rounds, substitutions were depleted in the buried core and around the modeled active sites and enriched on the solvent-exposed surface (Fig. 3f, Fig. S14ab), indicating that diversification concentrated on tolerant, exposed positions while the structural core and catalytic machinery were preserved. Relative to natural PKS sequences (ClusterCAD^62^), ∼45% of substitutions matched the position-specific consensus residue, a further ∼45% were to other residues common among natural PKS sequences at the aligned positions (≥5% frequency), with ∼9% to rare (<5%) and ∼1% to residues not observed at that position (Fig. 3g, Fig. S14). The substitutions were biochemically conservative, dominated by within-class exchanges (Fig. S14c) consistent with standard amino-acid substitution propensities (BLOSUM62; Fig. S14d).

Together, these analyses indicate that gLM2 distinguishes positions subject to strong catalytic constraints from regions that can tolerate diversification, while simultaneously modeling long-range structural and functional dependencies across the PKS assembly line.

### 2.6. High-performing PKS designs exhibit global sequence remodeling uncoupled from standard computational metrics

We next examined whether the functional performance of the generated PKSs could be explained by their global sequence relationships. Sequence similarity analysis showed that designs from both optimization rounds were substantially diverged from VL-Start and the CmiPKS M2 wildtype (WT) sequence and distributed across multiple branches of the respective trees (Fig. 4ab). High- and low-producing variants were interspersed throughout these branches, and neither the Round 1 nor Round 2 top-performing sequence belonged to a clade consistently enriched for high production. Functional improvement therefore did not result from convergence toward a single group of highly similar sequences.

Common computational measures of sequence and structural plausibility were also insufficient to identify high-performing designs. AlphaFold3 confidence scores did not positively correlate with VL production nor protein expression. Round 1 exhibited a weak negative association between pLDDT and titer, whereas no association was detected in Round 2 (Fig. S15, S16). Similarly, gLM2 pseudo-likelihood was not significantly associated with production in either Round 1 or Round 2 (Fig. S15, S16). These results indicate that high structural confidence and model likelihood are useful constraints on sequence plausibility but do not, by themselves, predict the expression or functional activity of a redesigned assembly line.

The final Round 2 top-performing PKS (R2-Top) produced up to 9.4-fold more valerolactam than VL-Start (Fig. 2d), yet differed from VL-Start at only 256 positions across the two redesigned regions (∼86% identity within the design windows). These substitutions were distributed throughout the two design windows rather than concentrated at the chimeric junctions or active site residues, and ranged from buried, conservative substitutions to more divergent surface-exposed changes (Fig. 4c). R2-Top substantially diverged from both wildtype parents of VL-Start. For the Round 1 design window, R2-Top aligned to wildtype FluPKS and CmiPKS at 64% and 82% identity respectively, in both cases at 99% query coverage. For the Round 2 design window, R2-Top aligned to FluPKS at 55% identity and CmiPKS at 81% identity, each at 100% query coverage.

**Figure 4.**
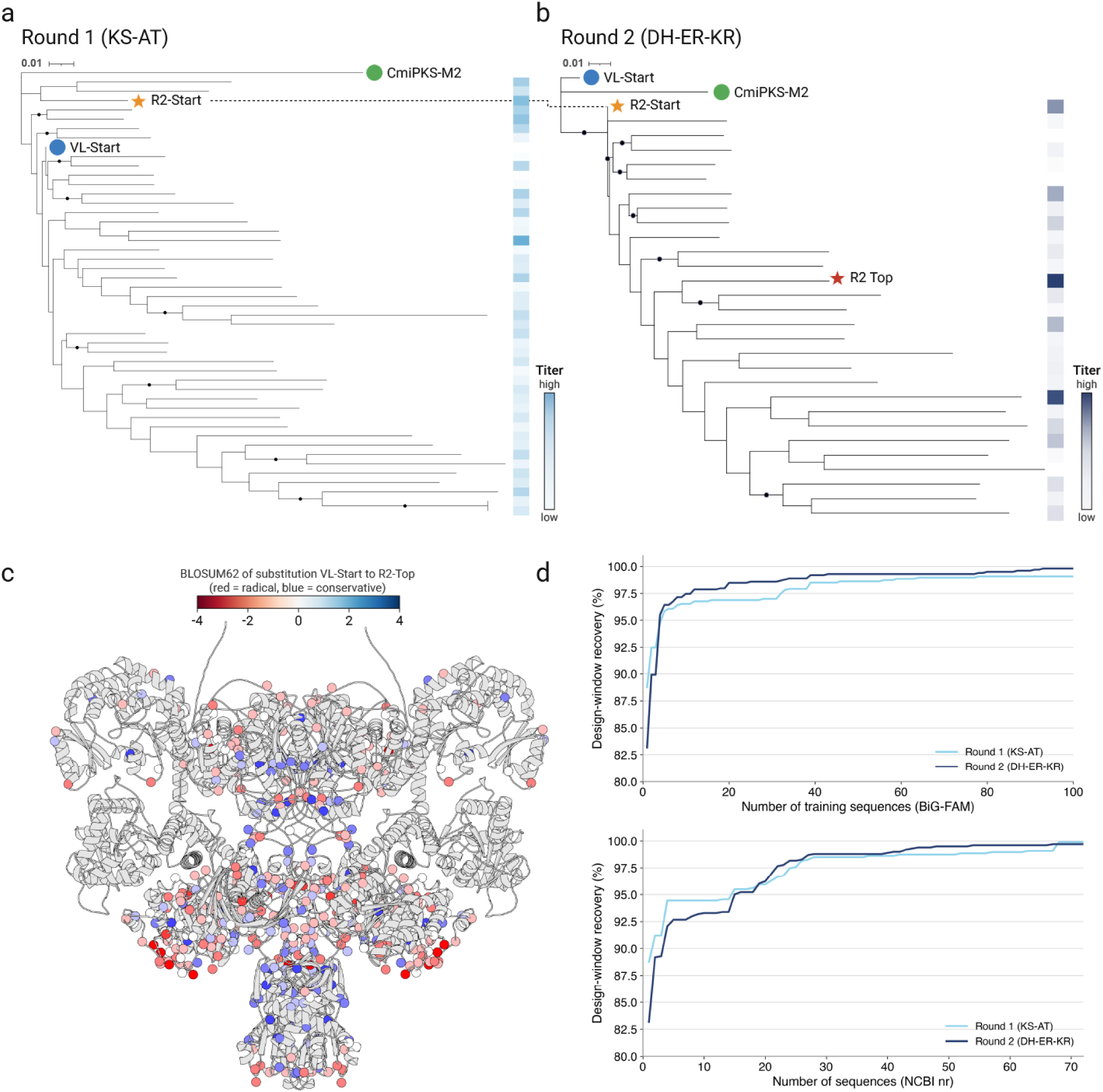
High-performing gLM2 designs are sequence-divergent and cannot be identified by global sequence or structure scores alone. a-b. Maximum-likelihood tree of experimentally tested Round 1 KS-AT **(a)** and Round 2 DH-ER-KR **(b)** designs, together with VL-Start and CmiPKS-M2 wildtype (WT). R2-Start and R2-Top are indicated; R2-Start corresponds to the top-performing Round 1 sequence used as the template for Round 2. Color strips indicate measured VL production with darker gradients corresponding to higher titers. Branches supported by bootstrap values >70% are marked with black circles. **c.** AlphaFold3-predicted structure of R2-Top with the 256 substituted residues relative to VL-Start shown as spheres and colored by BLOSUM62 substitution score (red = radical, blue = conservative). **d.** Cumulative recovery of the R2-Top Round 1 and Round 2 design windows as increasing numbers of matching sequences are combined from the BiG-FAM training set (top) or NCBI non-redundant (nr) database (bottom), illustrating that the optimized sequence is not reproduced by a single natural sequence. Figure created in BioRender.

Sequence searches further showed that R2-Top did not converge towards any individual natural PKS in either the BiG-FAM training set or NCBI non-redundant (nr) database. The closest match to both the Round 1 and Round 2 redesigned windows was CmiPKS M2, present in both databases. Recovering progressively larger fractions of the design windows required combining sequence segments from multiple natural examples, with up to 78 BiG-FAM sequences or 58 NCBI sequences required to reach 99% recovery (Fig. 4d). Therefore, the functional optimization of R2-Top stems from gLM2’s ability to interpolate across the PKS sequence space, integrating features from many distinct natural variants.

## 3. Discussion

Engineering biosynthetic assembly lines offer a route to programmable synthesis of complex molecules, and Type I PKSs are particularly attractive because their catalytic logic is modular and often encoded by domain order. Yet the difficulty of engineering PKSs through domain or module swaps suggests that the sequence constraints required for function are less modular than the chemistry itself. Consistent with this view, gLM2 designed variants with improved production contained substitutions distributed across catalytic domains, linkers, and positions far from the original chimeric boundary (Fig. 4c). However, our data do not establish that these distributed substitutions contribute equally, or even independently, to the observed gains in production. In particular, we cannot exclude the possibility that a small number of localized substitutions account for a substantial fraction of the improvement. Across the variants tested, product titer showed no clear relationship with substitution location, sequence similarity clustering, or global sequence and structural metrics (Fig. 4, Fig. S15, S16), preventing us from attributing improved function to specific substitutions or simple sequence-level properties. Resolving the causal contributions and epistatic interactions among substitutions will require systematic sampling of the large combinatorial sequence space of this assembly line, which may become increasingly feasible as long-DNA synthesis becomes less costly and the generative modeling framework developed here enables more targeted sampling of substitution combinations that are likely to preserve assembly-line function.

Our results motivate a different use of evolutionary information for PKS engineering. Evolution-guided approaches have largely used natural sequence diversity to identify recombination boundaries or compatible parts. Genomic language models can instead learn distributed constraints across natural systems and use them to generate new sequences in the context of a specified catalytic architecture. In our designs, these learned constraints enabled coordinated remodeling across large multi-domain regions rather than optimization of individual junctions or residues.

Notably, redesign of the complete DH-ER-KR region was performed without fixed residue-level constraints or a structural template. Sequence-only generation retained the features required for biosynthesis while improving product titer, suggesting genomic language models can successfully capture functional constraints contained in evolutionary sequence information. This capability is particularly relevant for genome-mined and evolutionarily divergent BGCs, where substrates, ligand-bound structures, and residue-level determinants of catalysis are often unknown.

Consistent with other generative protein and enzyme design studies, neither model likelihood nor predicted structural confidence tracked product titer ^63,64^, indicating that no single *in silico* metric captures the distributed requirements for assembly-line function. By learning long-range functional constraints directly from natural sequence diversity, gLMs provide a practical route to designing PKSs beyond naturally occurring sequence combinations. More broadly, this could enable biosynthetic assembly lines to be designed around desired sets of transformations, expanding the chemical space accessible through engineered biosynthesis.

## 5. Materials and Methods

### Enabling gLM2 generative design with discrete diffusion

The base gLM2 model (650M parameters)^2^ was originally trained using a masked language modeling objective with a fixed 30% random masking rate. To enable generative sequence design of multi-protein genomic contigs via discrete diffusion, we finetuned the model on the Open Genome (OG) dataset. Intergenic regions were removed, and training examples were constructed strictly from contiguous coding sequences (CDS) oriented in the same direction, with a context window of up to 4096 tokens. To mitigate training bias from overrepresented protein families, sample weights were applied based on MMseqs2^65^ clustering at 70% sequence identity. Specifically, each multi-protein contig was assigned a sample weight inversely proportional to the cluster size of its leading coding sequence.

The model was finetuned with a dynamic masking strategy, where the masking rate for each CDS was independently and uniformly sampled between 0% and 100%. A loss-reweighting objective was also applied, scaling the cross-entropy loss for each sequence by (1 − *p*), where *p* is the independently sampled masking probability. Training was performed for 1 epoch using a maximum sequence length of 4096 tokens and effective batch size of 128 examples. We utilized the AdamW optimizer^66^ with a peak learning rate of 4.0e-5, a 5000-step linear warmup, and cosine decay (Fig. S2). Training required 90 hours on a cluster of 8x H100 GPUs, employing bfloat16 half-precision and the FlashAttention^67^ library to optimize training efficiency.

### Finetuning gLM2 for biosynthetic gene cluster generation

To adapt the generative model for the programmable design of assembly lines, a second stage of finetuning was performed exclusively on the BiG-FAM database^27^ of biosynthetic gene clusters (BGCs) (accessed 01/2025). To enable domain-level programmability, the sequence tokenization scheme was augmented to include 1,999 antiSMASH^28^ HMM domain annotations. Each CDS was represented by its amino acid sequence preceded by its corresponding HMM domains, which were embedded as discrete special tokens sorted by sequence coordinates.

To accommodate the large scale of multi-protein functional complexes, the maximum context window was expanded to 16,384 tokens during finetuning. Training sequences longer than the context window were truncated. The model was finetuned for 4 epochs with effective batch size of 64 examples using the same uniform 0% to 100% masking strategy developed in Stage 1. Training utilized the AdamW optimizer with a peak learning rate of 4.0e-5, and a 5000-step warmup (Fig S2). Training required 80 hours on a cluster of 8x H100 GPUs. For generative sequence design and inference, we employed the Path Planning (P2) diffusion sampling method^68^ to iteratively decode masked tokens.

### Sequence redesign

A chimeric PKS sequence derived from FluPKS and CmiPKS (VL-Start)^24^ was used as a template for redesign with gLM2. The whole module redesign was divided into 2 rounds. Round 1 redesigned the KS and AT, and Round 2 redesigned the β-carbon processing enzymes (the reduction loop, RL) containing the DH, ER, and KR domains. Redesign boundaries were determined from an Alphafold3 predicted structure by identifying the N- and C-terminal limits of structurally well-packed catalytic domains. Round 1 designs were restricted to modifying residues 134 through 996, and round 2 designs were restricted to residues 1026 through 2006.

Sequences were generated using the fine-tuned gLM2 model by randomly masking residues within the design window, then iteratively unmasking residues with the P2 diffusion sampling method^68^ configured with temperature of 0.7 and an eta parameter of 0.1. For Round 1 designs, active site residues within the KS and AT domains were protected from mutation. The second round utilized the highest-producing variant from the first round as the template sequence and did not fix the active site residues. Following generation, all sequences were evaluated using the model’s pseudo-likelihood score and sorted in descending order. To eliminate low-complexity or degenerate sequences, a filter was applied by calculating the 1-gram Shannon entropy for each candidate sequence. Sequences with an entropy score falling below the 10th percentile of their respective generation pool were discarded. The Round 1 (KS-AT) experimental library consisted of 45 sequences and the Round 2 experimental library consisted of 28 designed sequences.

### Boltz-2 PKS ligand co-folding

Twenty-four ligand-bound PKS structures were retrieved from the RCSB PDB as PDBx/mmCIF (23 X-ray, 1.36–3.40 Å; one solution NMR ensemble, model 1), spanning the KS, AT, DH, KR, ACP and TE domains and including three multidomain constructs (KS-AT, KR-ER, DH-ER-KR). Structures were parsed programmatically with GEMMI 0.6.5^69^, and each was predicted with Boltz-2 from sequence and ligand identity without templates, pockets, contacts, or experimental coordinates. Each of the nine covalent complexes was run twice, unconstrained and with the observed bond supplied as a bond constraint. Predictions used three recycling steps and five diffusion samples per run on A40 GPUs with mmCIF output, with independent samples generated from separate random seeds. Predicted and experimental structures were superposed on protein Cα atoms; TM-score, pocket RMSD, symmetry-aware ligand heavy-atom RMSD and contact precision/recall/F1 computed in that frame. Because most entries predate the Boltz-2 training cutoff, these values quantify reconstruction of known complexes rather than generalization.

### Active site residue prediction

A VL-Start PKS model predicted by AlphaFold3^70^ was partitioned into its constituent catalytic domains (KS, AT, DH, ER, KR, ACP, TE) using residue boundaries (KS 1–550, AT 551–1000, DH 1001–1250, ER 1251–1500, KR 1501–1825, ACP 1826–2075, TE 2076–2488). The relevant domains were excised as monomers and co-folded with the phosphopantetheine-tethered substrate mimicking the intermediate it acts upon using Boltz-2. Residues were then reindexed to the parent full-length PKS numbering, and active-site residues were defined geometrically as any protein residue (ATOM records) with at least one heavy atom within 6 Å of any ligand atom (HETATM records), computed as all-atom Euclidean distances. Across three docking replicates, each hit was recorded then pooled for each domain-substrate pair and reduced to the union of unique positions.

### Tree-based sequence similarity analysis

For each design round, amino acid sequences from all generated PKS designs were aligned with MUSCLE using default parameters^71^. The corresponding CmiPKS M2 and VL-Start sequences were included as reference sequences. Wild-type FluPKS M1 was excluded from the tree calculation because its substantial sequence divergence from the designs resulted in poor alignment coverage and an exceptionally long branch. Maximum-likelihood trees were inferred from each alignment using IQ-TREE 3^72^ with default model-selection and tree-search settings and 1,000 bootstrap replicates. Trees were visualized with branch lengths representing substitutions per site with iToL^73^.

### Generated sequence analysis

To evaluate the structural and functional dependencies learned by the finetuned gLM2, a categorical Jacobian matrix was computed across the full template sequence. The top 0.01% long-range (≥100 residues apart) non-intra-domain Jacobian couplings were highlighted in Fig. 3a. We visualize the top 30 Jacobian couplings against AlphaFold3 predicted distances ≤ 12 Å both between KS-AT and AT-DH linker regions and between the DH-ER linker and KR domain.

To assess the diversity of the generated sequences, per-position Shannon entropy was calculated across generation samples (n=3,000 Round 1 designs and n=1,500 Round 2 designs). Native sequence recovery was defined as the fraction of samples perfectly matching the template sequence at the specified residue. Both sequence entropy and recovery were plotted using a 10 residue centered rolling mean.

Computational metrics of sequence plausibility were evaluated for their relationship to experimental titer. AlphaFold3 structural confidence (pLDDT) was computed as the mean whole-chain pLDDT across five models. Sequence pseudolikelihood was calculated as finetuned gLM2 masked language model pseudo-probability directly on the final experimentally tested sequences. The associations between these computational metrics and mean secretion titers were calculated using Spearman’s rank correlation (ρ).

Finally, to quantify the novelty of the generated regions versus natural sequences, cumulative sequence recovery was evaluated against the NCBI nr database via web BLAST^74^ and the model’s training set via BLASTp. Hits were evaluated in e-value rank order; a query position was counted as "recovered" if a hit aligned an identical residue at that position. Cumulative recovery curves were generated by calculating the fraction of the designed window covered by the union of identical matches from the top N contributing hits.

### Mutational landscape of designed substitutions

Substitutions were called per design against the VL-Start template by direct position-wise comparison, keeping positions within the design window (Round 1, 134-996; Round 2, 1026-2006; 10,103 total substitutions). Each template position was annotated on the AF3 predicted structure of the parent homodimer for burial using a backbone-only Cβ-neighbor count (binned into core/boundary/surface by Rosetta LayerSelector neighbor-count cutoffs), interface (dimer, cross-domain, and chimeric contacts), and active-site shell (see Active site residue prediction).

Enrichment by location was calculated as the observed fraction of substitutions in a class divided by its expected fraction, enrichment = (mutation events in class / total mutation events) / (designable positions in class / total designable positions in the window), where each design contributes one mutation event per mutated position and positions held fixed during generation (the Round 1 active-site residues) are excluded from the designable-position counts. 95% confidence intervals were computed by resampling the tested designs with replacement (1,000 iterations).

Substituted residues were scored against natural PKS variation from ClusterCAD module sequences aligned to the template (MAFFT --add --keeplength) and Henikoff-weighted, grading the identities at each position as consensus (most frequent), common (≥5%), rare (<5%), or unseen. A generic baseline reconstructed the change-conditional distribution from BLOSUM62 as *q*(*s*→*t*) ∝ *p_s_ p_t_* 2^(*S*(*s*,*t*)/2)^ where *p_s_*and *p_t_* are the background frequencies of residues *s* and *t*, and *S*(*s*,*t*) is the BLOSUM62 substitution score.

### Plasmid design and construction

PKS DNA fragments were codon-optimized using BaseBuddy^75^ (match codon usage, *Pseudemonas putida* TaxID: 160488) and synthesized (Twist). Phanta Max (Vazyme) Super-Fidelity DNA Polymerase was used to amplify DNA and plasmid fragments for plasmid construction. Golden Gate assembly with TypeIIS enzyme BsmbI-v2 (NEBridge Golden Gate Assembly Kit) was used to construct the plasmids. The mixture was transformed into NEB Stable competent *E. coli*. Sequence verification was achieved via a combination of 1) isolation using Qiaprep Spin Miniprep kit (Qiagen) and Whole Plasmid Nanopore sequencing (Angstrom Innovation) and 2) colony PCR followed by LevSeq^76^ (long read <u>e</u>very <u>v</u>ariant <u>seq</u>uencing) (Plasmidsaurus).

### *P. putida* Strain Construction

The parent strain used in this study was taken from the original Lee study^24^: *P. putida* strain LM3 harboring a single copy of the Flv helper pathway at the RV *attb* site. An additional copy of the FlvJ enzyme was integrated using the SAGE integration system^50^ at the TG1 *attb* site. All PKS designs were integrated using a SAGE integration system. Competent cells were prepared for integration with the following steps. An overnight culture of LM3-FlvJ was washed three times with an equal volume of 10% glycerol resuspended in 1/4th original volume. Each electroporation used 60uL of prepared cells. Each PKS plasmid (a serine integrase *attP* site plasmid containing the PKS expression cassette) and a non-replicating helper integrase expression plasmid (60 ng each) were introduced into the competent cells through electroporation at 2.5 kV using a BTX electroporation system. After electroporation the cells recovered in 0.9 mL of LB medium at 30 °C for 2 hrs. Finally, recovered cells were plated on LB agar plates with the appropriate antibiotic selection marker for overnight growth. Successful integrations were verified via colony PCR amplification of *the recombination junction*.

## Culture conditions

For valerolactam production, seed cultures inoculated from glycerol stocks into selective LB and grown for 16 hours. Fresh production medium was prepared composed of 1X premixed M9 salts (Teknova), 5 g/L yeast extract (Gibco Bacto), 20 g/L glucose (MP Biomedicals), 5mM L-aspartic acid (Sigma-Aldrich), and 1X trace metals solution (Teknova). Cultures were set up in 48-well plates with 1.5ml volume and 1% inoculation, and grown in a 1000 rpm orbital shaker incubator at 30C for 96 hours. After initial screening of both libraries, production culture medium was optimized by supplementation with 1X Grace’s amino acid mix (Sunrise Science).

## Valerolactam measurements

Following cultivation, valerolactam was best extracted from cultures by cell lysis at -80° C for ≥ 4 hours, mixing 1:3 volume of media:methanol, followed by shaking at 225 rpm at 37° C for 4 hours. Plates were centrifuged at 5000 *x*g for 90 minutes, and 100 uL of supernatant was pipetted into plates for LCMS analysis. Standards of δ-valerolactam were purchased from MilliporeSigma. LCMS was conducted on an Agilent Ultivo Triple Quadrupole LC/MS with a Kintex C18 analytical column (Phenomenex, 100x3.0 mm). The mobile phase was composed of water (Milli-Q ultrapure water purification system) and methanol (Sigma-Aldrich), both with 0.1% formic acid (Sigma-Aldrich). The run method defined a 5uL sample injection, with an organic phase gradient ranging from 5-95% over 11 minutes at 40C. Response data was collected using ESI positive ion mode with scans for mass range *m*/*z* 90–220 and SIM *m*/*z* 100. Collected data was analyzed using Agilent MassHunter software.

## Protein Expression Measurements

Following cultivation, 1mL of culture was pelleted via centrifugation at 5000g for 5 minutes, followed by three successive rounds of washes with 1mL of 1X PBS (Teknova) before final pelleting and storage at -80C. Protein was extracted from cell pellets and tryptic peptides were prepared by following established proteomic sample preparation protocol^77^). Briefly, cell pellets were resuspended in Qiagen P2 Lysis Buffer (Qiagen, Germany) to promote cell lysis. Proteins were precipitated with addition of 1 mM NaCl and 4 x vol acetone, followed by two additional washes with 80% acetone in water. The recovered protein pellet was homogenized by pipetting mixing with 100 mM ammonium bicarbonate in 20% methanol. Protein concentration was determined by the DC protein assay (BioRad, USA). Protein reduction was accomplished using 5 mM tris 2-(carboxyethyl)phosphine (TCEP) for 30 min at room temperature, and alkylation was performed with 10 mM iodoacetamide (IAM; final concentration) for 30 min at room temperature in the dark. Overnight digestion with trypsin was accomplished with a 1:50 trypsin:total protein ratio. The resulting peptide samples were analyzed on an Agilent 1290 UHPLC system coupled to a Thermo Scientific Orbitrap Exploris 480 mass spectrometer for discovery proteomics^78^. Briefly, peptide samples were loaded onto an Ascentis® ES-C18 Column (Sigma–Aldrich, USA) and were eluted from the column by using a 10 minute gradient from 98% solvent A (0.1 % FA in H2O) and 2% solvent B (0.1% FA in ACN) to 65% solvent A and 35% solvent B. Eluting peptides were introduced to the mass spectrometer operating in positive-ion mode and were measured in data-independent acquisition (DIA) mode with a duty cycle of 3 survey scans from m/z 380 to m/z 985 and 45 Tandem mass spectrometry (MS2) scans with precursor isolation width of 13.5 m/z to cover the mass range. DIA raw data files were analyzed by an integrated software suite DIA-NN^79^. The database used in the DIA-NN search (library-free mode) is *P. putida* (TaxID: 160488) latest Uniprot proteome FASTA sequences plus the protein sequences of the heterologous proteins and common proteomic contaminants. DIA-NN determines mass tolerances automatically based on first pass analysis of the samples with automated determination of optimal mass accuracies. The retention time extraction window was determined individually for all MS runs analyzed via the automated optimization procedure implemented in DIA-NN. Protein inference was enabled, and the quantification strategy was set to Robust LC = High Accuracy. Output main DIA-NN reports were filtered with a global false discovery rate set at 0.01 (FDR ≤ 0.01) on both the precursor level and protein group level. The Top3 method, which is the average MS signal response of the three most intense tryptic peptides of each identified protein, was used to plot the quantity of the targeted proteins in the samples^80,81^.

## Acknowledgements

The authors thank Isaac Donnell and Namil Lee for their valuable discussions and support throughout this work, as well as *P. putida* for its patience with our repeated genetic interventions.

## Author Contributions

Conceptualization: KEI, NL, YH, AC, TK, JDK

Methodology: KEI, NL, YH, AC, JDK

Investigation: NL, KEI, YH, AC, JDK

Data curation: YH, AC

Validation: led by NL, KEI

Plasmid/strain construction and verification: NL, KEI, MH, VG, AJ, JA, JB

Strain cultivation, lysis, and VL extraction: NL, KEI, MH, VG, AJ, JA

LC-MS methods development and analysis: KEI, NL, JB

Proteomic analysis: JWG, YC, CJP

Formal analysis: NL, KEI, YH, AC

Resource: YH, AC, TK, JDK

Software: YH, AC

Project Administration: YH, AC, TK, JDK

Visualization: NL, KEI, YH, AC

Funding acquisition: YH, AC, TK, JDK

Supervision: YH, AC, TK, JDK

Writing-original draft: KEI, NL, YH, AC

Writing-review & editing: All authors

## Data and code availability

The OpenGenome (OG) dataset used for finetuning is available on HuggingFace at https://huggingface.co/datasets/tattabio/OG. The finetuned gLM2 model is available at https://huggingface.co/tattabio/gLM2_650M_bgc_decoder. Model code is available at https://github.com/TattaBio/gLM2_decoder. Designed sequences are found in Extended Data 1.

The generated mass spectrometry proteomics data have been deposited to the ProteomeXchange Consortium via the PRIDE partner repository with the dataset identifier PXD083871^82^. DIA-NN is freely available for download from https://github.com/vdemichev/DiaNN.

## References

1. Hwang, Y., Cornman, A. L., Kellogg, E. H., Ovchinnikov, S. & Girguis, P. R. Genomic language model predicts protein co-regulation and function. Nat. Commun. 15, 2880 (2024).

2. Cornman, A., et al. The OMG dataset: An Open MetaGenomic corpus for mixed-modality genomic language modeling. in The Thirteenth International Conference on Learning Representations (2024).

3. Nguyen, E. et al. Sequence modeling and design from molecular to genome scale with Evo. Science 386, eado9336 (2024).

4. Brixi, G. et al. Genome modelling and design across all domains of life with Evo 2. Nature 652, 1349–1361 (2026).

5. Merchant, A. T., King, S. H., Nguyen, E. & Hie, B. L. Semantic design of functional de novo genes from a genomic language model. Nature 649, 749–758 (2026).

6. King, S. H. et al. Generative design of bacteriophages with genome language models. Science 393, eaec2657 (2026).

7. Nivina, A., Yuet, K. P., Hsu, J. & Khosla, C. Evolution and diversity of assembly-line polyketide synthases: Focus review. Chem. Rev. 119, 12524–12547 (2019).

8. Davies, J. Specialized microbial metabolites: functions and origins. J. Antibiot. (Tokyo) 66, 361–364 (2013).

9. Newman, D. J. & Cragg, G. M. Natural products as sources of new drugs over the nearly four decades from 01/1981 to 09/2019. J. Nat. Prod. 83, 770–803 (2020).

10. Nielsen, J. & Keasling, J. D. Engineering cellular metabolism. Cell 164, 1185–1197 (2016).

11. Yin, K. et al. Polyketide synthase-based controlled synthesis of polycyclopropanated fuel molecules. Nat. Commun. 17, 6904 (2026).

12. Wang, Z. et al. Engineered polyketide synthases enable a microbial chassis for recyclable plastics with tunable properties. Nat. Biotechnol. 1–10 (2026).

13. Hagen, A. et al. Engineering a polyketide synthase for in vitro production of adipic acid. ACS Synth. Biol. 5, 21–27 (2016).

14. Jenke-Kodama, H., Börner, T. & Dittmann, E. Natural biocombinatorics in the polyketide synthase genes of the actinobacterium Streptomyces avermitilis. PLoS Comput. Biol. 2, e132 (2006).

15. Nivina, A., Herrera Paredes, S., Fraser, H. B. & Khosla, C. GRINS: Genetic elements that recode assembly-line polyketide synthases and accelerate their diversification. Proc. Natl. Acad. Sci. U. S. A. 118, e2100751118 (2021).

16. Sherman, D. H. The Lego-ization of polyketide biosynthesis. Nat. Biotechnol. 23, 1083–1084 (2005).

17. Bozhüyük, K. A. J. et al. Evolution-inspired engineering of nonribosomal peptide synthetases. Science 383, eadg4320 (2024).

18. Mabesoone, M. F. J. et al. Evolution-guided engineering of trans-acyltransferase polyketide synthases. Science 383, 1312–1317 (2024).

19. Klaus, M. et al. Protein-protein interactions, not substrate recognition, dominate the turnover of chimeric assembly line polyketide synthases. J. Biol. Chem. 291, 16404–16415 (2016).

20. Kalkreuter, E. et al. Computationally-guided exchange of substrate selectivity motifs in a modular polyketide synthase acyltransferase. Nat. Commun. 12, 2193 (2021).

21. Donadio, S., Staver, M. J., McAlpine, J. B., Swanson, S. J. & Katz, L. Modular organization of genes required for complex polyketide biosynthesis. Science 252, 675–679 (1991).

22. Xu, Y. et al. Metabolic engineering of Escherichia coli for polyamides monomer δ-valerolactam production from feedstock lysine. Appl. Microbiol. Biotechnol. 104, 9965–9977 (2020).

23. von Tiedemann, P., Anwar, S., Kemmer-Jonas, U., Asadi, K. & Frey, H. Synthesis and solution processing of nylon-5 ferroelectric thin films: The renaissance of odd-nylons? Macromol. Chem. Phys. 221, 1900468 (2020).

24. Lee, N. et al. Retrobiosynthesis of unnatural lactams via reprogrammed polyketide synthase. Nat. Catal. 8, 389–402 (2025).

25. Cornman, A., Tranzillo, M., Zulaybar, N. G., Bouzit, I. & Hwang, Y. Linear-time prediction of proteome-scale microbial protein interactions. Proc. Natl. Acad. Sci. U. S. A. 123, (2026).

26. Austin, J., Johnson, D. D., Ho, J., Tarlow, D. & van den Berg, R. Structured denoising diffusion models in discrete state-spaces. arXiv [cs.LG] (2021) doi:10.48550/arXiv.2107.03006.

27. Kautsar, S. A., Blin, K., Shaw, S., Weber, T. & Medema, M. H. BiG-FAM: the biosynthetic gene cluster families database. Nucleic Acids Res. 49, D490–D497 (2021).

28. Blin, K., Shaw, S., Medema, M. H. & Weber, T. The antiSMASH database version 5. Nucleic Acids Res. 54, D522–D526 (2026).

29. Miyanaga, A. et al. Identification of the Fluvirucin B2 (Sch 38518) Biosynthetic Gene Cluster from Actinomadura fulva subsp. indica ATCC 53714: substrate Specificity of the β-Amino Acid Selective Adenylating Enzyme FlvN. Biosci. Biotechnol. Biochem. 80, 935–941 (2016).

30. Amagai, K., Takaku, R., Kudo, F. & Eguchi, T. A unique amino transfer mechanism for constructing the β-amino fatty acid starter unit in the biosynthesis of the macrolactam antibiotic cremimycin. Chembiochem 14, 1998–2006 (2013).

31. Gay, D. C. et al. A close look at a ketosynthase from a trans-acyltransferase modular polyketide synthase. Structure 22, 444–451 (2014).

32. Tang, Y., Kim, C.-Y., Mathews, I. I., Cane, D. E. & Khosla, C. The 2.7-Å crystal structure of a 194-kDa homodimeric fragment of the 6-deoxyerythronolide B synthase. Proc. Natl. Acad. Sci. U. S. A. 103, 11124–11129 (2006).

33. Tang, Y., Chen, A. Y., Kim, C.-Y., Cane, D. E. & Khosla, C. Structural and mechanistic analysis of protein interactions in module 3 of the 6-deoxyerythronolide B synthase. Chem. Biol. 14, 931–943 (2007).

34. Miyazawa, T., Hirsch, M., Zhang, Z. & Keatinge-Clay, A. T. An in vitro platform for engineering and harnessing modular polyketide synthases. Nat. Commun. 11, 80 (2020).

35. Zhang, L. et al. Characterization of Giant Modular PKSs Provides Insight into Genetic Mechanism for Structural Diversification of Aminopolyol Polyketides. Angew. Chem. Int. Ed Engl. 56, 1740–1745 (2017).

36. Keatinge-Clay, A. T. Polyketide synthase modules redefined. Angew. Chem. Int. Ed Engl. 56, 4658–4660 (2017).

37. Miyazawa, T., Fitzgerald, B. J. & Keatinge-Clay, A. T. Preparative production of an enantiomeric pair by engineered polyketide synthases. Chem. Commun. (Camb.) 57, 8762–8765 (2021).

38. Huang, Z. et al. Plug-and-play engineering of modular polyketide synthases. Nat. Chem. Biol. 1–7 (2025).

39. Dutta, S. et al. Structure of a modular polyketide synthase. Nature 510, 512–517 (2014).

40. Ostrowski, M. P., Cane, D. E. & Khosla, C. Recognition of acyl carrier proteins by ketoreductases in assembly line polyketide synthases. J. Antibiot. (Tokyo) 69, 507–510 (2016).

41. Kapur, S., Chen, A. Y., Cane, D. E. & Khosla, C. Molecular recognition between ketosynthase and acyl carrier protein domains of the 6-deoxyerythronolide B synthase. Proc. Natl. Acad. Sci. U. S. A. 107, 22066–22071 (2010).

42. Klaus, M. & Grininger, M. Engineering strategies for rational polyketide synthase design. Nat. Prod. Rep. 35, 1070–1081 (2018).

43. Whicher, J. R. et al. Structural rearrangements of a polyketide synthase module during its catalytic cycle. Nature 510, 560–564 (2014).

44. Khosla, C., Herschlag, D., Cane, D. E. & Walsh, C. T. Assembly Line Polyketide Synthases: Mechanistic Insights and Unsolved Problems. Biochemistry 53, 2875–2883 (2014).

45. McCullough, T. M. et al. Structure of a modular polyketide synthase reducing region. Structure 31, 1109–1120.e3 (2023).

46. Wong, F. T., Chen, A. Y., Cane, D. E. & Khosla, C. Protein-protein recognition between acyltransferases and acyl carrier proteins in multimodular polyketide synthases. Biochemistry 49, 95–102 (2010).

47. Robbins, T., Liu, Y.-C., Cane, D. E. & Khosla, C. Structure and mechanism of assembly line polyketide synthases. Curr. Opin. Struct. Biol. 41, 10–18 (12/2016).

48. Hirsch, M., Fitzgerald, B. J. & Keatinge-Clay, A. T. How cis-acyltransferase assembly-line ketosynthases gatekeep for processed polyketide intermediates. ACS Chem. Biol. 16, 2515–2526 (2021).

49. Passaro, S., et al. Boltz-2: Towards accurate and efficient binding affinity prediction. bioRxivorg (2025) doi:10.1101/2025.06.14.659707.

50. Elmore, J. R. et al. High-throughput genetic engineering of nonmodel and undomesticated bacteria via iterative site-specific genome integration. Sci. Adv. 9, eade1285 (2023).

51. Wrenbeck, E. E., Azouz, L. R. & Whitehead, T. A. Single-mutation fitness landscapes for an enzyme on multiple substrates reveal specificity is globally encoded. Nat. Commun. 8, 15695 (2017).

52. Zargar, A. et al. Chemoinformatic-guided engineering of polyketide synthases. J. Am. Chem. Soc. 142, 9896–9901 (2020).

53. Barajas, J. F. et al. Structural insights into dehydratase substrate selection for the borrelidin and fluvirucin polyketide synthases. J. Ind. Microbiol. Biotechnol. 46, 1225–1235 (2019).

54. Tsai, S.-C. S. & Ames, B. D. Structural enzymology of polyketide synthases. Methods Enzymol. 459, 17–47 (2009).

55. Akey, D. L. et al. Crystal structures of dehydratase domains from the curacin polyketide biosynthetic pathway. Structure 18, 94–105 (2010).

56. Keatinge-Clay, A. T. & Stroud, R. M. The structure of a ketoreductase determines the organization of the beta-carbon processing enzymes of modular polyketide synthases. Structure 14, 737–748 (2006).

57. Zheng, J., Gay, D. C., Demeler, B., White, M. A. & Keatinge-Clay, A. T. Divergence of multimodular polyketide synthases revealed by a didomain structure. Nat. Chem. Biol. 8, 615–621 (2012).

58. Kwan, D. H. & Leadlay, P. F. Mutagenesis of a modular polyketide synthase enoylreductase domain reveals insights into catalysis and stereospecificity. ACS Chem. Biol. 5, 829–838 (2010).

59. Zhang, Z. et al. Protein language models learn evolutionary statistics of interacting sequence motifs. Proc. Natl. Acad. Sci. U. S. A. 121, e2406285121 (2024).

60. Bagde, S. R., Mathews, I. I., Fromme, J. C. & Kim, C.-Y. Modular polyketide synthase contains two reaction chambers that operate asynchronously. Science 374, 723–729 (2021).

61. Yuzawa, S. et al. Comprehensive in vitro analysis of acyltransferase domain exchanges in modular polyketide synthases and its application for short-chain ketone production. ACS Synth. Biol. 6, 139–147 (2017).

62. Eng, C. H. et al. ClusterCAD: a computational platform for type I modular polyketide synthase design. Nucleic Acids Res. 46, D509–D515 (2018).

63. Johnson, S. R. et al. Computational scoring and experimental evaluation of enzymes generated by neural networks. Nat. Biotechnol. 43, 396–405 (2025).

64. Braun, M. et al. Computational enzyme design by catalytic motif scaffolding. Nature 649, 237–245 (2026).

65. Steinegger, M. & Söding, J. MMseqs2 enables sensitive protein sequence searching for the analysis of massive data sets. Nat. Biotechnol. 35, 1026–1028 (2017).

66. Loshchilov, I. & Hutter, F. Decoupled weight decay regularization. arXiv [cs.LG] (2017) doi:10.48550/arXiv.1711.05101.

67. Shah, J., et al. FlashAttention-3: Fast and accurate attention with asynchrony and low-precision. arXiv [cs.LG] (2024) doi:10.48550/arXiv.2407.08608.

68. Peng, F. Z. et al. Path Planning for masked diffusion model sampling. arXiv [cs.LG] (2025) doi:10.48550/arXiv.2502.03540.

69. Wojdyr, M. GEMMI: A library for structural biology. J. Open Source Softw. 7, 4200 (2022).

70. Abramson, J. et al. Accurate structure prediction of biomolecular interactions with AlphaFold 3. Nature 630, 493–500 (2024).

71. Edgar, R. C. MUSCLE: multiple sequence alignment with high accuracy and high throughput. Nucleic Acids Res. 32, 1792–1797 (2004).

72. Nguyen, L.-T., Schmidt, H. A., von Haeseler, A. & Minh, B. Q. IQ-TREE: a fast and effective stochastic algorithm for estimating maximum-likelihood phylogenies. Mol. Biol. Evol. 32, 268–274 (2015).

73. Letunic, I. & Bork, P. Interactive Tree Of Life (iTOL): an online tool for phylogenetic tree display and annotation. Bioinformatics 23, 127–128 (2007).

74. Altschul, S. F., Gish, W., Miller, W., Myers, E. W. & Lipman, D. J. Basic local alignment search tool. J. Mol. Biol. 215, 403–410 (1990).

75. Schmidt, M. et al. Maximizing heterologous expression of engineered type I polyketide synthases: Investigating Codon optimization strategies. ACS Synth. Biol. 12, 3366–3380 (2023).

76. Long, Y. et al. LevSeq: Rapid Generation of Sequence-Function Data for Directed Evolution and Machine Learning. ACS Synth. Biol. 14, 230–238 (2025).

77. Chen, Y. Alkaline-SDS cell lysis of microbes with acetone protein precipitation for proteomic sample preparation in 96-well plate format v1. (2021) doi:10.17504/protocols.io.6qpvr6xjpvmk/v1.

78. Chen, Y., Gin, J. & J Petzold, C. Discovery proteomic (DIA) LC-MS/MS data acquisition and analysis v2. (2022) doi:10.17504/protocols.io.e6nvwk1z7vmk/v2.

79. Demichev, V., Messner, C. B., Vernardis, S. I., Lilley, K. S. & Ralser, M. DIA-NN: neural networks and interference correction enable deep proteome coverage in high throughput. Nat. Methods 17, 41–44 (2020).

80. Ahrné, E., Molzahn, L., Glatter, T. & Schmidt, A. Critical assessment of proteome-wide label-free absolute abundance estimation strategies. Proteomics 13, 2567–2578 (2013).

81. Silva, J. C., Gorenstein, M. V., Li, G.-Z., Vissers, J. P. C. & Geromanos, S. J. Absolute quantification of proteins by LCMSE: a virtue of parallel MS acquisition. Mol. Cell. Proteomics 5, 144–156 (2006).

82. Perez-Riverol, Y. et al. The PRIDE database resources in 2022: a hub for mass spectrometry-based proteomics evidences. Nucleic Acids Res. 50, D543–D552 (2022).

