## Supplementary Information for "Generative Design of New-to-nature Biosynthetic Assembly Lines with Genomic Language Modeling"

* These authors contributed equally

Affiliations:

### Supplemental Notes 1-3

**Supplementary Note 1. Optimization of strain engineering between Round 1 and Round 2**

The first production conducted (Fig. S9a) used the exact strain reported by Lee *et al.* as cmipks163 which was generated with a distinct integration vector (pBH026, PKS cargo, BxbI *attP*, *lacI*) that differs from the pJB integration system (pJB035, PKS cargo, co-expressed BxbI integrase, BxbI *attP*). We therefore integrated the cmipks163 sequence in the pJB system with nucleotide level modifications to remove the TypeIIS recognition sequence for BsmbI (used for Golden Gate assembly of design fragments) into the same base strain used for the Round 1 designs.

During these design campaigns, Klass *et al*[83]. identified genomic instability at the integration sites within the same landing pad in AG5577 (Fig. S7)[83]. In brief, when multiple adjacent sites are occupied, the genome can undergo spontaneous homologous recombination with long term use that removes the integrated cargo, a potential issue with the Round 1 library (Fig. S7, Round 1 Strains) for long term use. Because *Pseudomonas putida* carries many copies of its genome, these deleterious HDR events are difficult to detect with the standard PCR-based verification protocol. In order to avoid this issue entirely, Round 2 designs were integrated into the R4 position in SAGE Landing Pad 2 (Fig. S7, Round 2 Strains) using a modified pJB vector that lacks the *cis*-element of the R4 integrase. Therefore, the Round 1 and Round 2 designs are not ideally comparable due to uncharacterized individual effects of the 1) different integration locus and 2) co-expression of the BxbI integrase in R1. R2-Start was integrated with both systems and proteomic analysis indicated increased titer as a function of expression (Fig. S12d), though other factors likely contribute to the discrepancy.

**Supplementary Note 2. Active site and catalytic residue preservation in unconstrained generation**

Predicted active-site residues (n = 118 within the DH-ER-KR design window) were defined as residues with any heavy atom within 6 Å of the docked substrate in the per-domain cofolding models (Methods). Within this set we annotated the specific catalytic residues of the three reductive-loop domains: the DH His-Asp dyad (H1054/D1224)[54,55,84], the KR short-chain dehydrogenase/reductase (SDR) catalytic tetrad (K1944, S1968, Y1981, N1985)[54,56,85], and the ER Lys/Asp pair (K1743/D1767)[57,58,86]. The DH catalytic histidine and the KR catalytic tyrosine were identified from their conserved sequence motifs (DH: His-box HxxxGxxxxP motif; KR: YxxxN) and confirmed to lie within the 6 Å ligand-contact shell in the cofolding models. The remaining catalytic residues were assigned by structural proximity to them, and the ER Lys/Asp pair by structural alignment to the SpnER2 enoylreductase (PDB 3SLK)[57]. We note that the DH His-box of cmipks163 lacks the proline of the canonical HxxxGxxxxP motif (**H**RVL**G**RVIV**S**), so the H1054/D1224 dyad was assigned from its geometry and ligand contact rather than the sequence motif alone[54,55,84]. The ER stereochemistry "fingerprint" tyrosine, which sets product configuration but is not catalytic[87], was excluded.

Across all 1,500 unconstrained gLM2 samples, the DH and KR catalytic residues were retained in ≥99.8% of sequences, and the full active-site set showed a 38-fold lower median substitution rate than the remainder of the design window (Fig. S11). The only appreciable variation among catalytic residues was at ER D1767 (98% conserved). Modular PKS ER domains lack a single classic catalytic nucleophile, its lysine contributes most to catalysis, and no single mutation abolishes activity[57,58,86]. The catalytically dominant ER Lys1743 remained 99.9% conserved, and none of the 28 synthesized Round 2 designs mutated any catalytic residue.

**Supplementary Note 3. Validating gLM2 categorical-Jacobian couplings in experimentally determined PKS structures**

Because gLM2 and AlphaFold3 are trained on overlapping evolutionary data, agreement between a Categorical Jacobian coupling and an AlphaFold3 contact could reflect a shared learned prior rather than physical reality. We therefore validated the couplings against experimentally determined structures with no structure predictor in the loop, computing the categorical Jacobian directly on the full-length parent ORFs of three deposited PKS structures (DEBS2/EryAII, PDB 2QO3[32]; the juvenimicin PKS subunit, PDB 8G7W[45]; and Lsd14, PDB 7S6B[60]), aligning each sequence to its deposited chain(s), and measuring every long-range ($|i-j| > 50$), non-intra-domain coupled pair directly in the deposited coordinates. Of the strongest such couplings ($|J| > 4$), 97% (102 of 105) are inter-domain Cβ-Cβ distances within 12 Å in the DEBS KS-AT didomain, 94% (79 of 84) in the juvenimicin reduction loop, and 93% (270 of 290) in the intact Lsd14 module (Fig. S15a-c). These couplings concentrate at interfaces of established importance: the post-AT linker packing against the KS-AT linker (resolved in the DEBS didomain and the intact Lsd14 module and whose boundary governs functional AT-domain exchange[61]) and the DH-ER linker (KRs subdomain) and its packing against the catalytic KR domain ($KR_C$) as critical to the reduction loop interfaces[45].

For VL-Start, mapping the couplings onto its AlphaFold3 model showed that 94% of the top-scoring long-range inter-domain pairs are within 12 Å Cβ-Cβ distance in the model (Fig. S15d). Not every strong coupling corresponds to a distance within this threshold: the residues surrounding the phosphopantetheine attachment sites of the loading ACP ($ACP_L$) and the downstream ACP are strongly coupled yet lie 71-86 Å apart in the model (Fig. 3c), consistent with a dependency that reflects ACP sequence and functional similarity rather than structural proximity.

#### Supplemental Figures 1-16

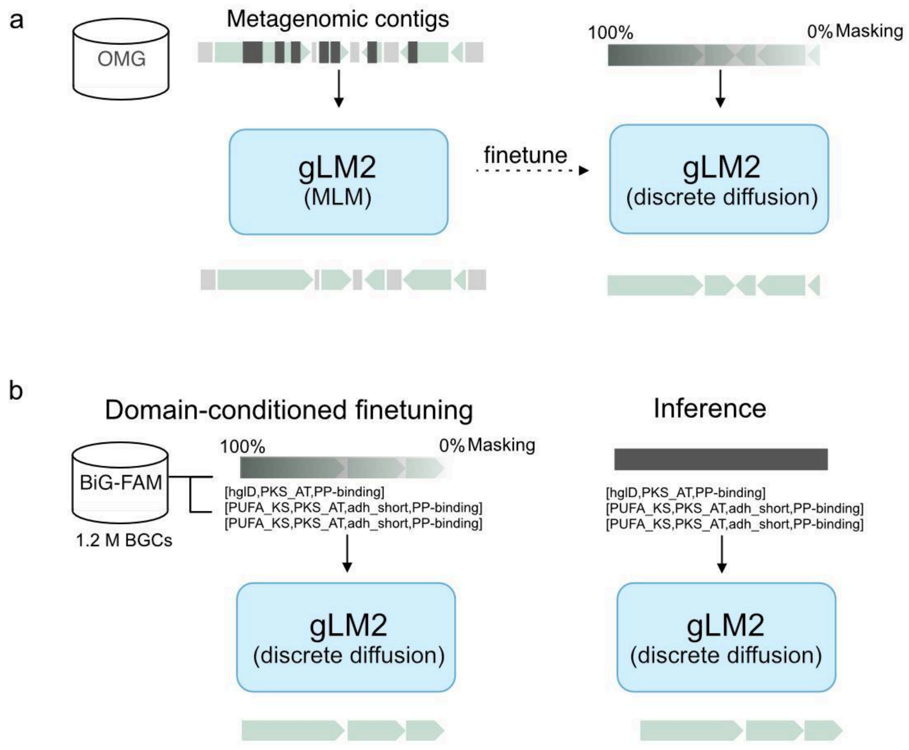

**Figure S1. Two-stage finetuning framework for generative sequence design and domain-level conditioning. a.** In Stage 1, gLM2 is finetuned on multi-protein OpenGenome contigs (with intergenic regions removed) via discrete diffusion with dynamic masking (0-100%), enabling multi-gene sequence generation and in-painting. **b.** In Stage 2, the model is adapted to BiG-FAM BGCs using an expanded vocabulary of 1,999 antiSMASH domain tokens. These special tokens are prepended to coding sequences, enabling domain-level conditional generation.

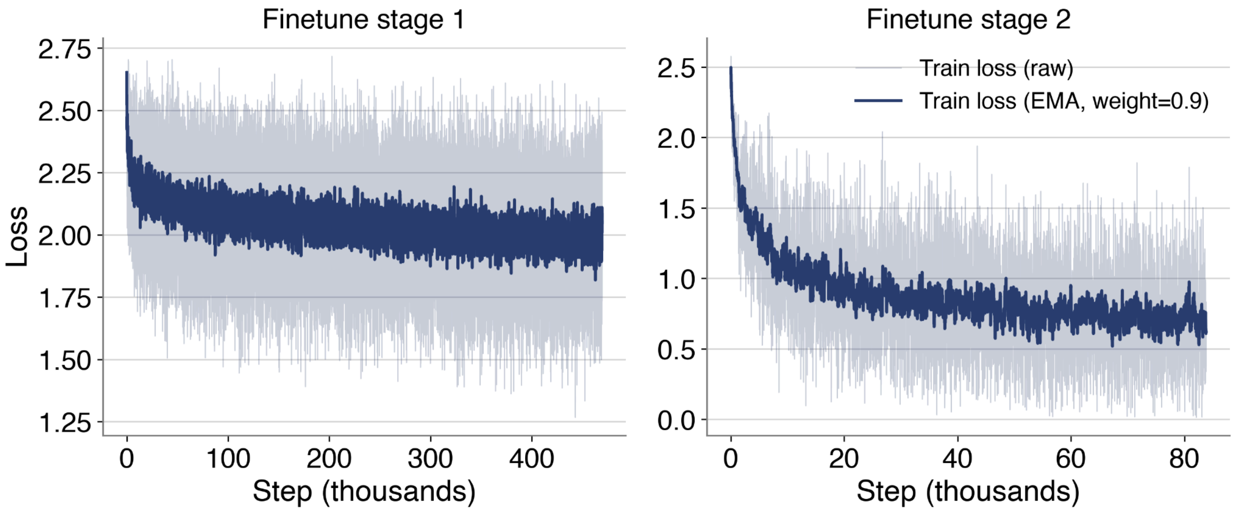

**Figure S2. Training loss curves for the two fine-tuning stages.** Per-step training loss (light) with exponential moving average smoothing (dark, weight = 0.9) for stage 1 finetuning on OpenGenome (left) and stage 2 finetuning on BiG-FAM (right).

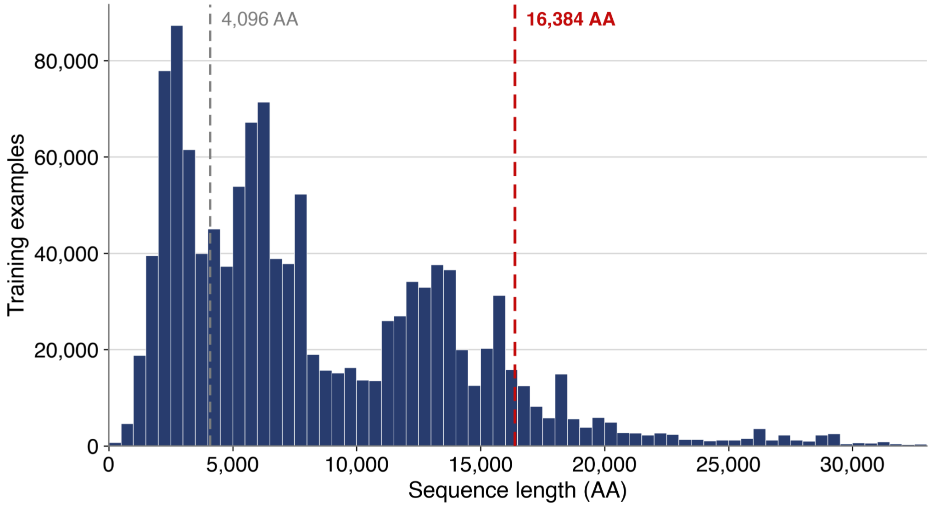

**Figure S3. Training sequence length distribution.** Distribution of training example lengths in amino acids (AA) for the dataset (median 6,591 AA; 95th percentile 18,688 AA). The dashed grey line indicates the base gLM2 context window of 4,096 tokens. The dashed red line indicates the extended window used for gLM2 fine tuning. Training examples with length beyond the extended context window were truncated.

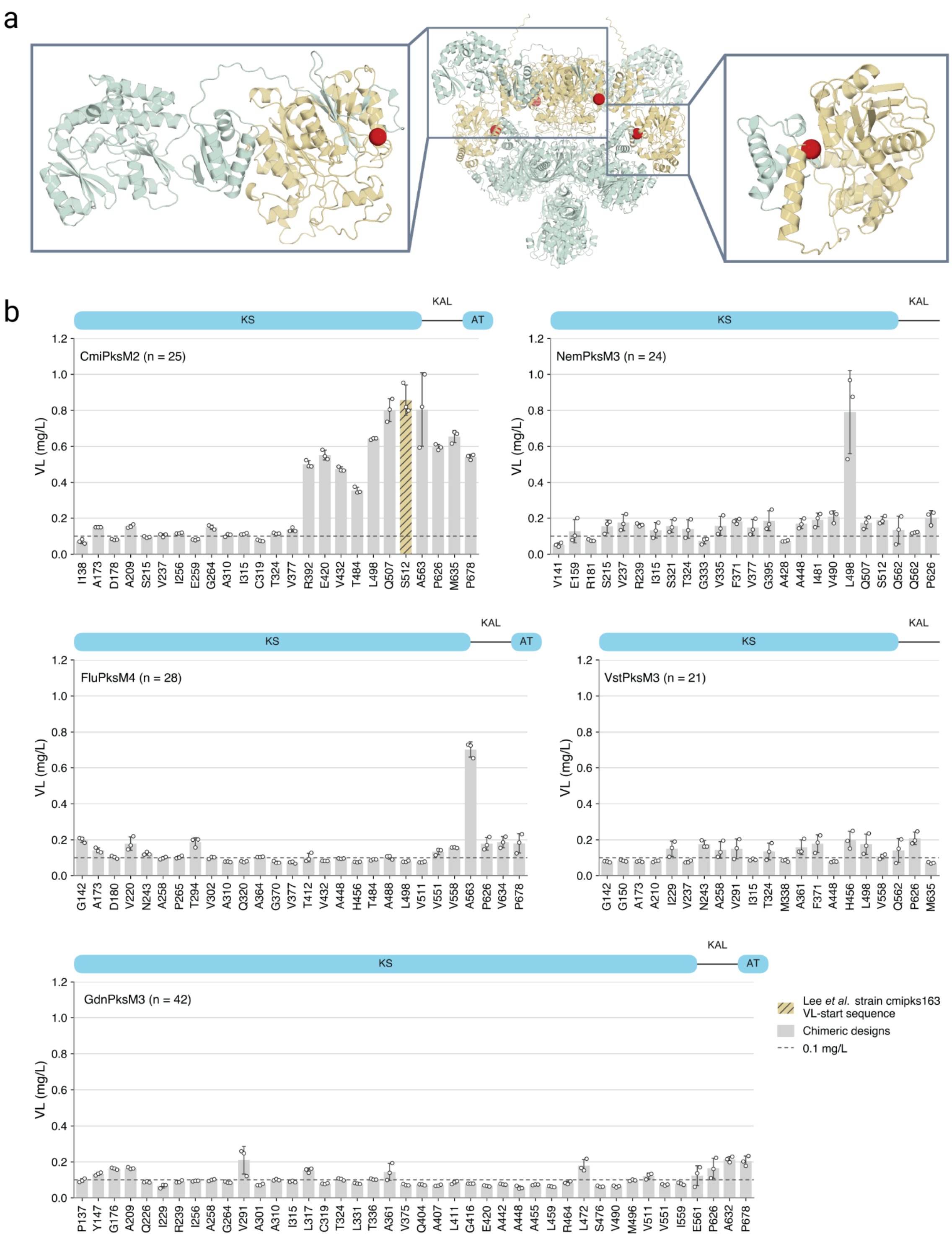

**Figure S4. Selection of template sequence for gLM2-based optimization from Lee *et al.*[24] a.** VL-Start is chimerized at the KS adapter (KSad) near the end of the KS domain between FluPKS M1 (tan) and CmiPKS M2 (teal). Chimera junctions shown as red spheres. **b.** VL-Start was selected as the top producing variant in a 140 member library of PKS sequences chimerized at various conserved positions across the KS domain[24]. Data shown was not generated as a part of this study. See Lee *et al.* Supplementary Figure 18.

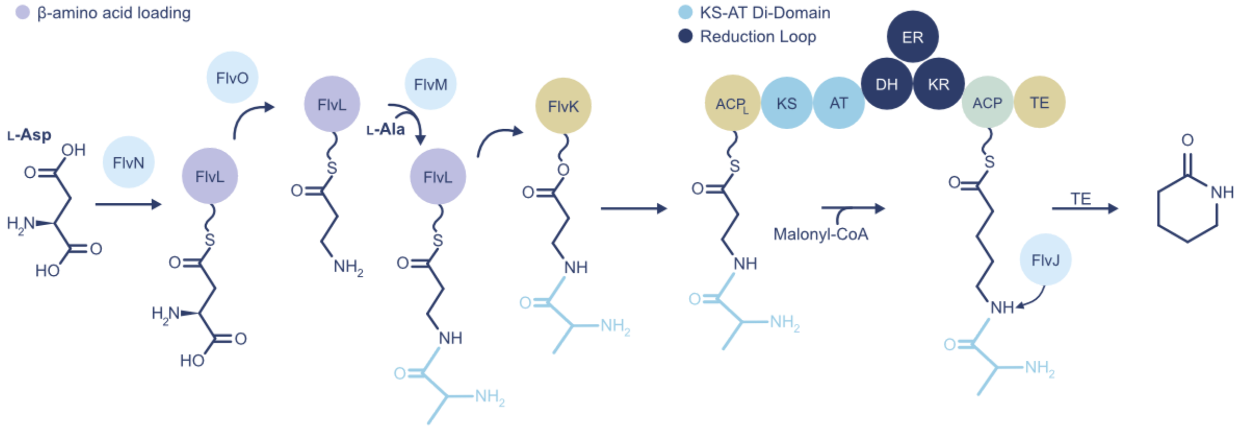

**Figure S5. Biosynthesis of δ-valerolactam (VL).** A cascade of monofunctional enzymes from the fluvirucin biosynthetic pathway prepare the aminoacyl starting unit by 1) loading of L-Asp onto ACP FlvJ by FlvN, 2) decarboxylation of L-Asp by FlvO, and 3) FlvM-mediated adenylation with L-Ala. The modified amino acid chain is transferred onto FlvK, which coordinates loading onto the first PKS domain, $ACP_L$, to initiate VL biosynthesis. $ACP_L$ translocates the substrate onto the KS while the AT loads extender unit malonyl-CoA onto the downstream ACP. Holo-KS and ACP domains coordinate for a decarboxylative Claisen condensation, transferring the growing polyketide onto the downstream ACP, which then shuttles the substrate to each successive reduction domain for full reduction of the β-carbon. Accessory enzyme FlvJ deaminates the processed substrate, exposing a primary amine and enabling TE-based hydrolysis and cyclization into the free lactam.

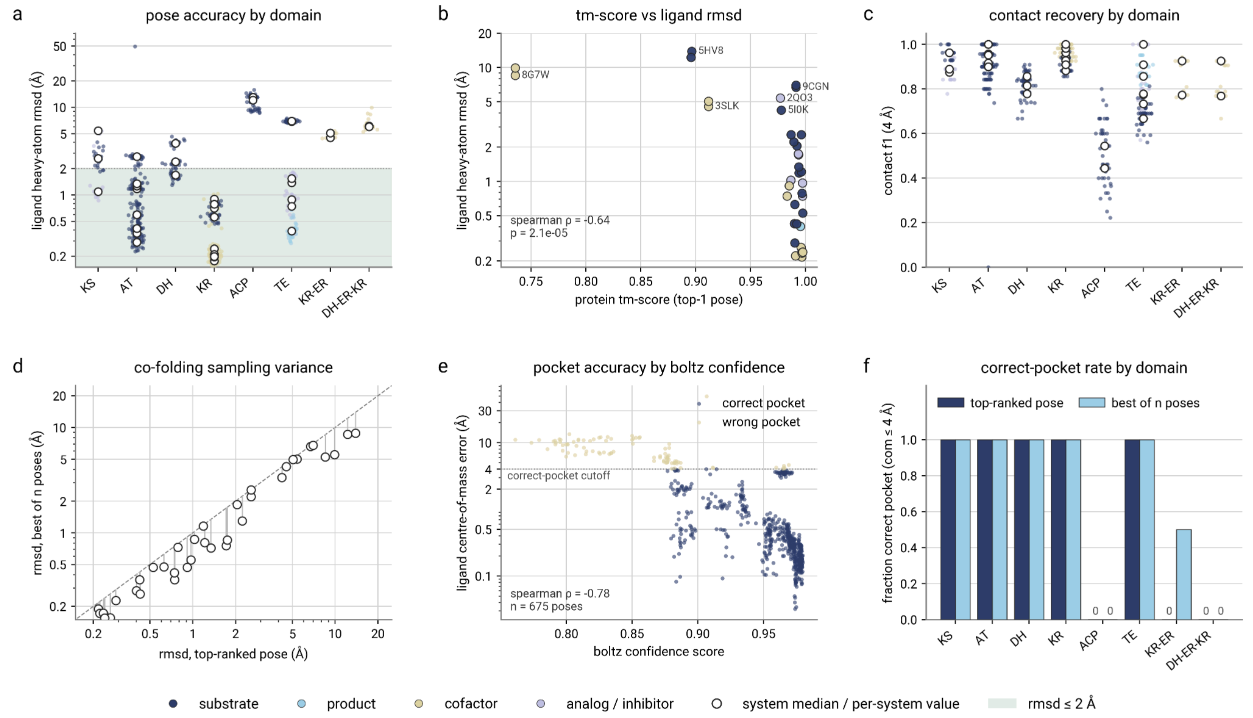

**Figure S6. Boltz-2 co-folding accuracy on ligand-bound PKS structures.** All panels summarize 675 scored poses from 33 prediction jobs over 24 experimental structures. Ligand metrics were computed after superposing the predicted protein onto the experimental chain on Cα atoms alone. RMSD is symmetry-aware and restricted to ligand atoms resolved in the deposition. Point color denotes ligand class. **a.** RMSD by domain poses, per-system medians, and the 2Å band. **b.** RMSD vs TM-score, outliers labelled, $\rho = -0.64$, $n = 37$. **c.** Contact F1 at 4Å by domain. **d.** Best-of-N vs top-1 RMSD. **e.** COM error vs confidence for all 675 poses, colored by correct-pocket, $\rho = -0.78$. **f.** Correct-pocket fraction, top-1 vs best-of-N.

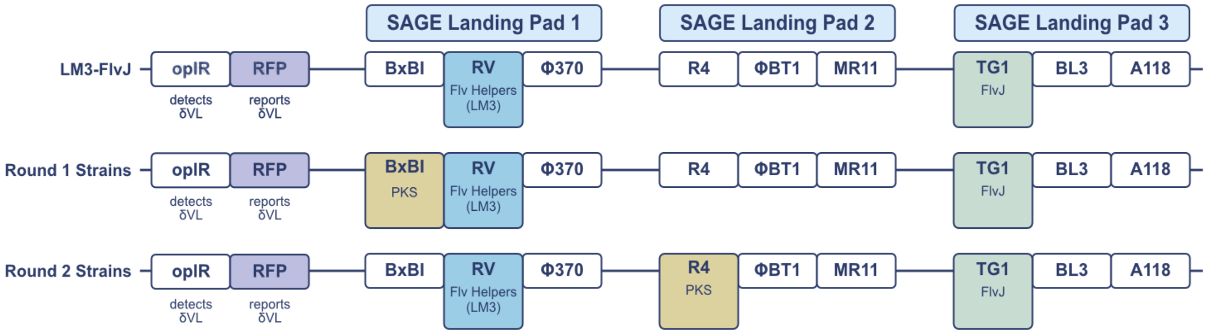

**Figure S7. SAGE integration map for base and production strains**. LM3-FlvJ is derived from Guss Lab *P. putida* strain AG5577 with three landing pads each containing three serine integrase sites to insert engineered constructs. This strain contains knockouts (ΔPP_5182, ΔDavAB) and a previously described valerolactam biosensor (ΔoplBA::sens2). The RV site contains the Flv helper pathway shown in Fig. S5 with an additional copy of FlvJ integrated at the TG1 site. Round 1 strains integrate the PKS at the BxBI site. Round 2 strains integrate the PKS at the distal landing pad (R4 site) for increased genomic stability.

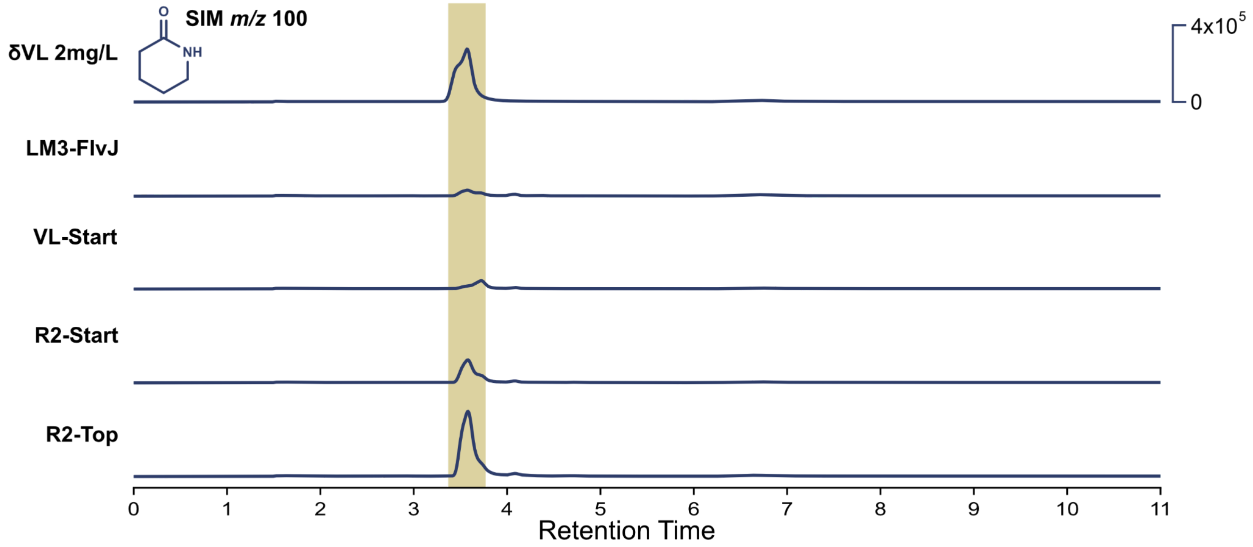

**Figure S8. LCMS trace for δ-valerolactam standard and strains.** Strains shown include LM3-FlvJ (base strain), VL-Start, and best performing gLM2 redesigns by round (R2-Start and R2-Top). Peaks shown represent peak extraction using SIM in positive mode at *m/z* 100.

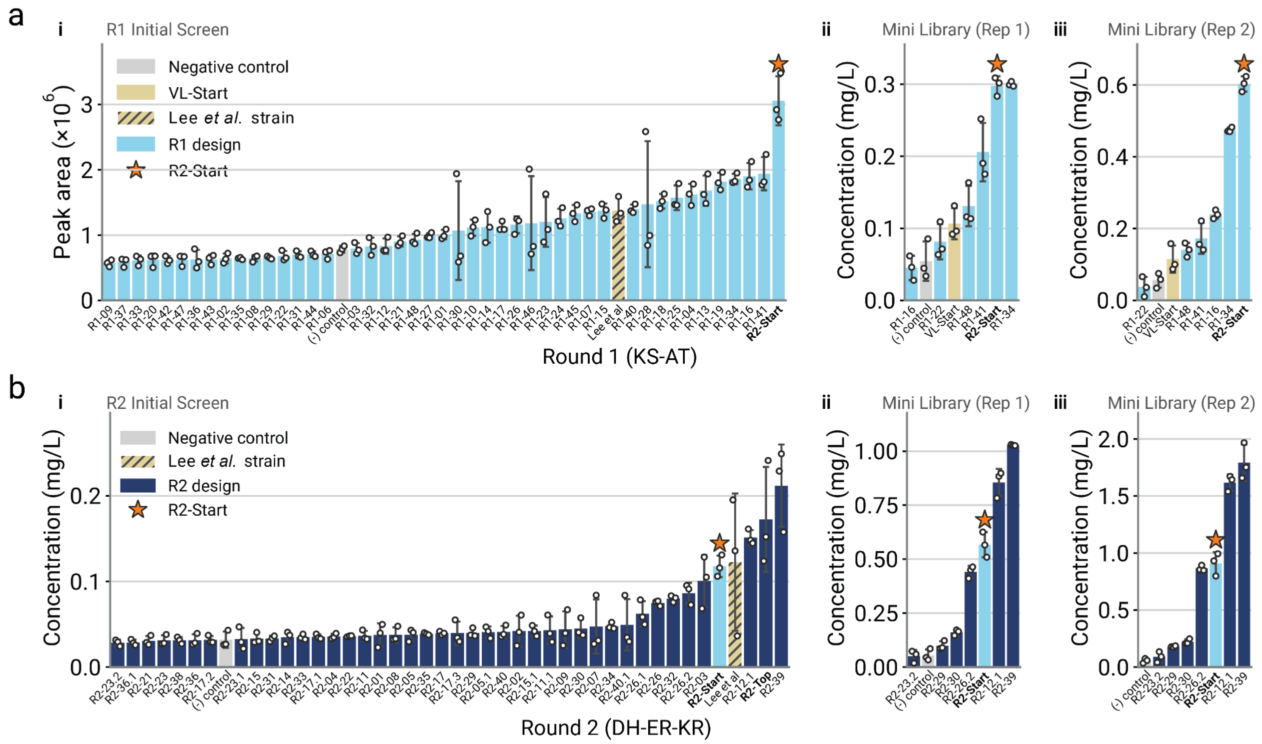

**Figure S9**. **Independent cultivations of all Round 1 and Round 2 strains.** Production titer for all assayed **(a)** Round 1 and **(b)** Round 2 designs and cloning-derived mutants from independent cultivations. **i.** Initial screen of each design round with the published strain from Lee *et al*. **ii.** Independent cultivation of selected designs (mini library) with corrected positive control strain (cmipks163 sequence integrated with the pJB035 system, Supplementary Note 1). **iii.** Repeat cultivation of mini library supplemented with Grace's amino acid mix and optimized freeze-thaw lysis (Methods). Individual points are technical replicates (n = 3), bars are sorted ascending (mean ± SD).

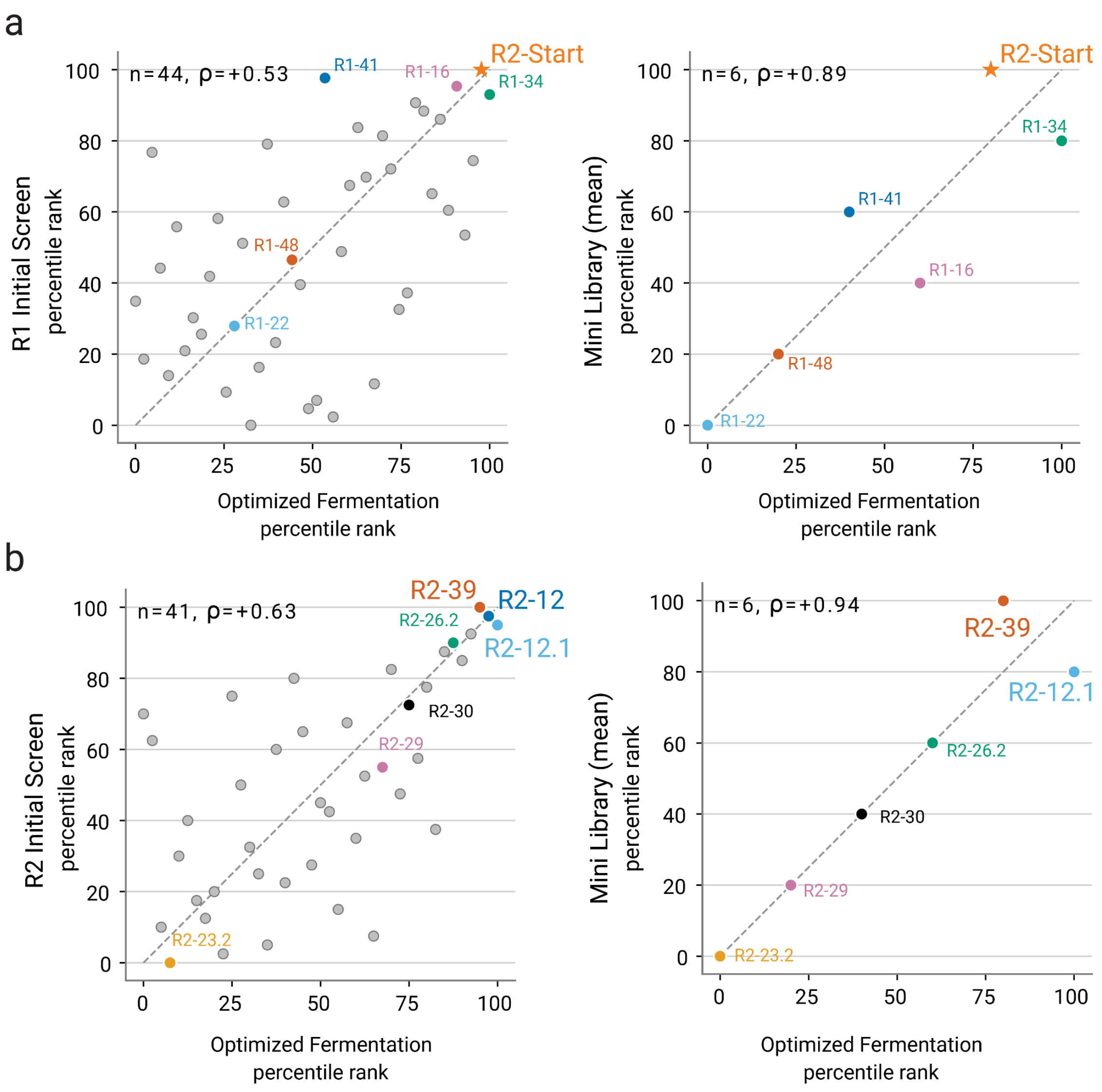

**Figure S10. Rank order of top designs is consistent over independent cultivations.** Percentile rank of each strain represented in multiple independent cultivations. Left: Initial screens of **(a)** Round 1 and **(b)** Round 2 are in agreement with results of the final optimized cultivation in which the medium was supplemented with Grace's amino acid mix and optimized extraction via freeze-thaw (n=44, rho=0.53 and n=41, rho=0.63 for Round 1 and Round 2, respectively). Right: Mean percentile rank order of both cultivations of the mini library strongly agrees with the optimized cultivation (n=6, rho=0.89 and 0.94 for Round 1 and Round 2, respectively). R2-12 = R2-Top, R2-12.1 is a cloning-derived mutant of R2-12 (A1374T, A1466V).

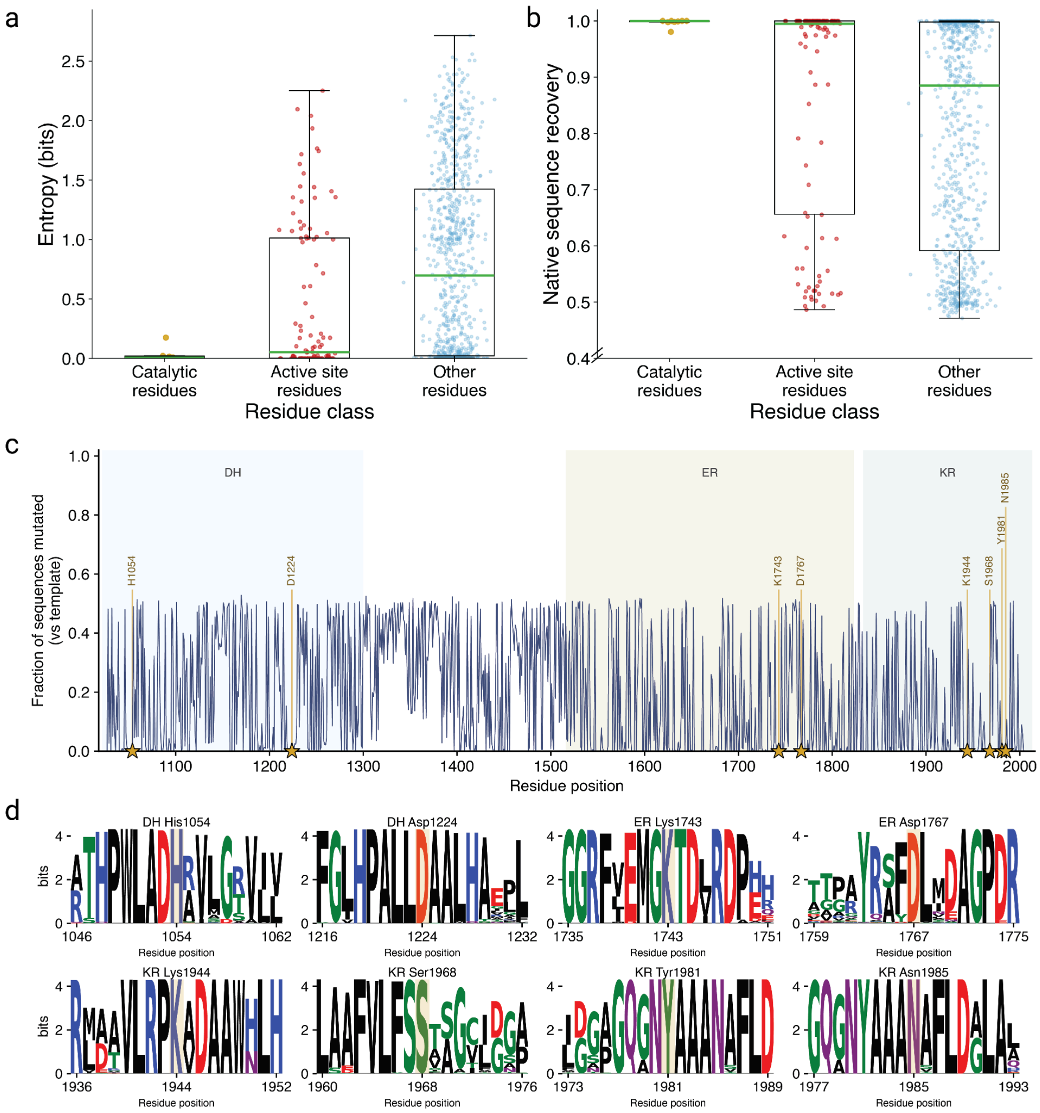

**Figure S11. Round 2 (DH-ER-KR) generation conserves catalytic residues and preferentially retains native active site residues** (for catalytic and active site residue determination, see Methods). None of the 28 synthesized Round 2 designs contained any mutated catalytic residue. Per-position Shannon entropy **(a)** and native sequence recovery **(b)** across all 1,500 unconstrained Round 2 generation samples (500 each at mask rates 0.3/0.5/0.7), at catalytic residues (gold, n = 8; labelled in **(c-d)**, the remaining active site residues (n = 110) determined by the Boltz-2 cofolding pipeline, and the remainder of the design region (blue, n = 863). Median entropy was 0.03 bits across all 118 active site positions versus 0.70 bits elsewhere (Mann-Whitney $P = 1.7\times10^{-8}$); median recovery was 1.00 versus 0.89 (Mann-Whitney $P = 2.6\times10^{-6}$). The catalytic residues were the most strongly conserved class (median recovery 1.00, entropy ≈ 0 bits). **c.** Per-position fraction of the 1,500 samples differing from the Round 2 starting sequence across the DH, ER, and KR domains. The eight catalytic residues (gold; DH H1054/D1224 dyad, ER K1743/D1767 pair, KR K1944/S1968/Y1981/N1985 tetrad; stars below the axis) are mutated in only 1.9% of samples and are unchanged in all tested designs. **d.** Sequence logos centred on each catalytic residue (gold highlight). Each catalytic position is essentially invariant while its flanking positions vary, indicating the conservation is residue-specific rather than a locally conserved patch. The single more-variable catalytic residue, ER D1767 (98% conserved), is the less-critical half of the ER reductase's Lys/Asp pair, which unlike the DH and KR active sites has no lone catalytic nucleophile and no single residue whose mutation abolishes activity[57,58]; the key ER K1743 remains 99.9% conserved.

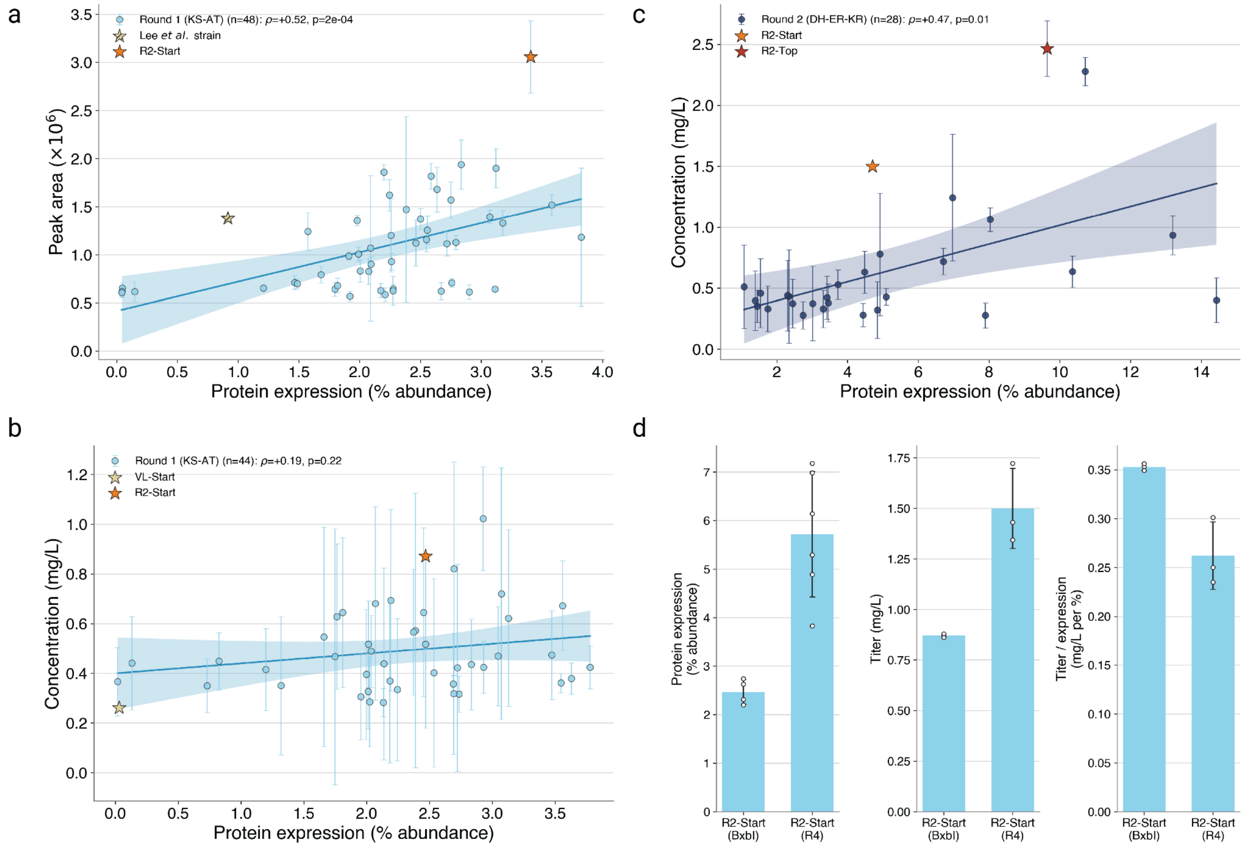

**Figure S12. Proteomic analysis of designed PKS sequences. a-b.** Proteomic-based analysis of PKS expression vs VL production titer of Round 1 designs from 2 independent experiments. Correlation in **b)** is weak due to higher variance of measured titer resulting from the large scale cultivation. **c.** Expression of Round 2 PKS designs correlate with higher titer, but do not explain high expressing, low producing designs. Notably, the top two producing designs' titers cannot be explained fully by increased expression. **d.** Expression differences between R2-Start in Round 1 (BxbI *attB* integration + constitutive co-expression of BxbI integrase) and in Round 2 (R4 *attB* integration, no co-expression of R4 recombinase).

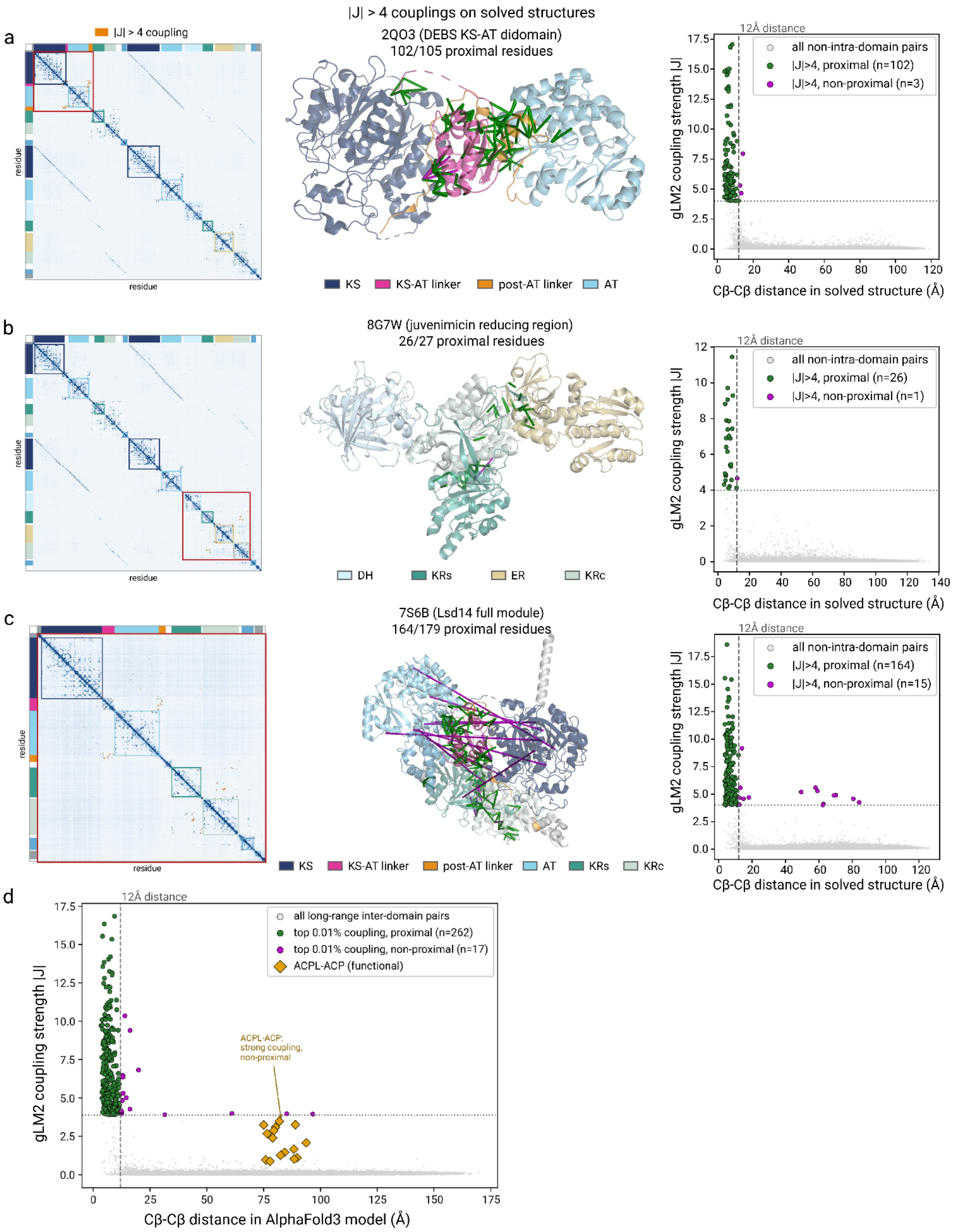

**Figure S13. gLM2 couplings correspond to close distances in experimentally determined PKS structures.** **a-c.** The categorical Jacobian computed directly on the full-length parent sequence of each deposited PKS structure. Left: the full-ORF Jacobian, colored by domain (key below each structure); the red box marks the region resolved in the deposited structure. Center: the deposited structure with the strongest long-range non-intra-domain couplings (|J| > 4) drawn as Cβ-Cβ rods (green: distances <12Å, fuchsia: distances >12Å). Right: coupling strength (|J|) versus the Cβ-Cβ distance measured in the deposited structure for every long-range non-intra-domain pair; the |J| > 4 couplings are overwhelmingly within 12 Å Cβ-Cβ distances (**a**, 102/105 = 97%; **b**, 26/27 = 96%; **c**, 164/179 = 92%; dashed line, 12Å). **d.** Coupling strength versus Cβ-Cβ distance in the AF3 model of VL-Start for all top-scoring long-range inter-domain pairs; 94% of the top 0.01% are within a <12 Å Cβ-Cβ distance. The $ACP_L$-ACP phosphopantetheine pairs (Fig. 3c) are the labelled non-proximal outliers. Per-pair values in Extended Data 2.

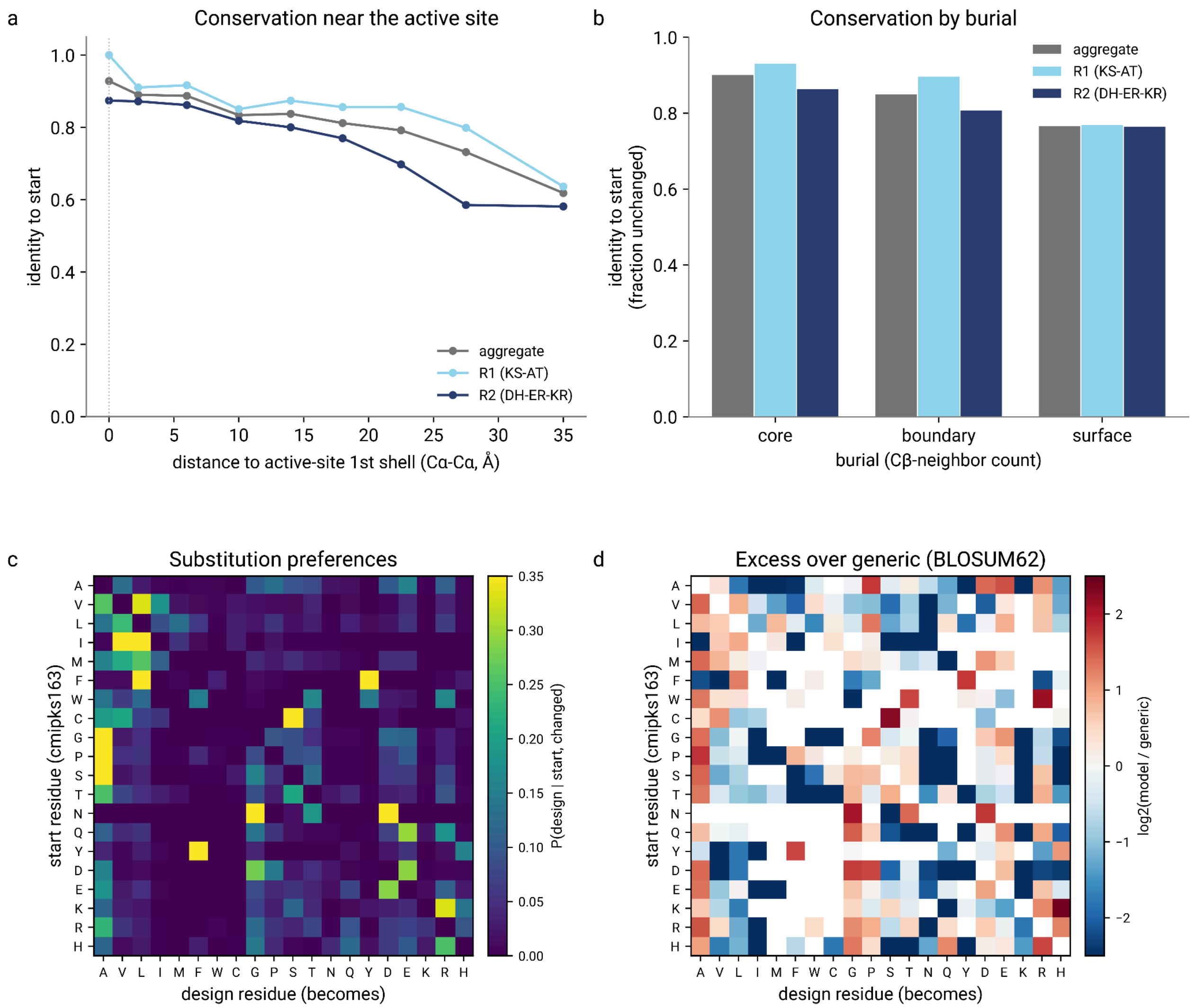

**Figure S14. Extended mutational landscape of the tested chimeric-PKS design library. a.** Sequence identity to the template versus distance to the nearest modeled active-site (first-shell) residue (Cα–Cα). **b.** Sequence identity to the starting template by burial class (Rosetta Cβ-neighbor count), for the aggregate library and each design round. **c.** Change-conditional amino-acid substitution matrix, P(design residue | starting residue, conditional on a change), aggregated over the library. **d.** Excess of the model's substitution preferences over a generic BLOSUM62 expectation, $\log_2$(model ÷ generic); the model's preferences track generic exchangeability (Spearman $\rho = 0.61$) but are modestly more conservative (52% of substitutions to a BLOSUM-favorable target vs 46% expected). Analyses use the tested designs (10,103 in-window substitutions).

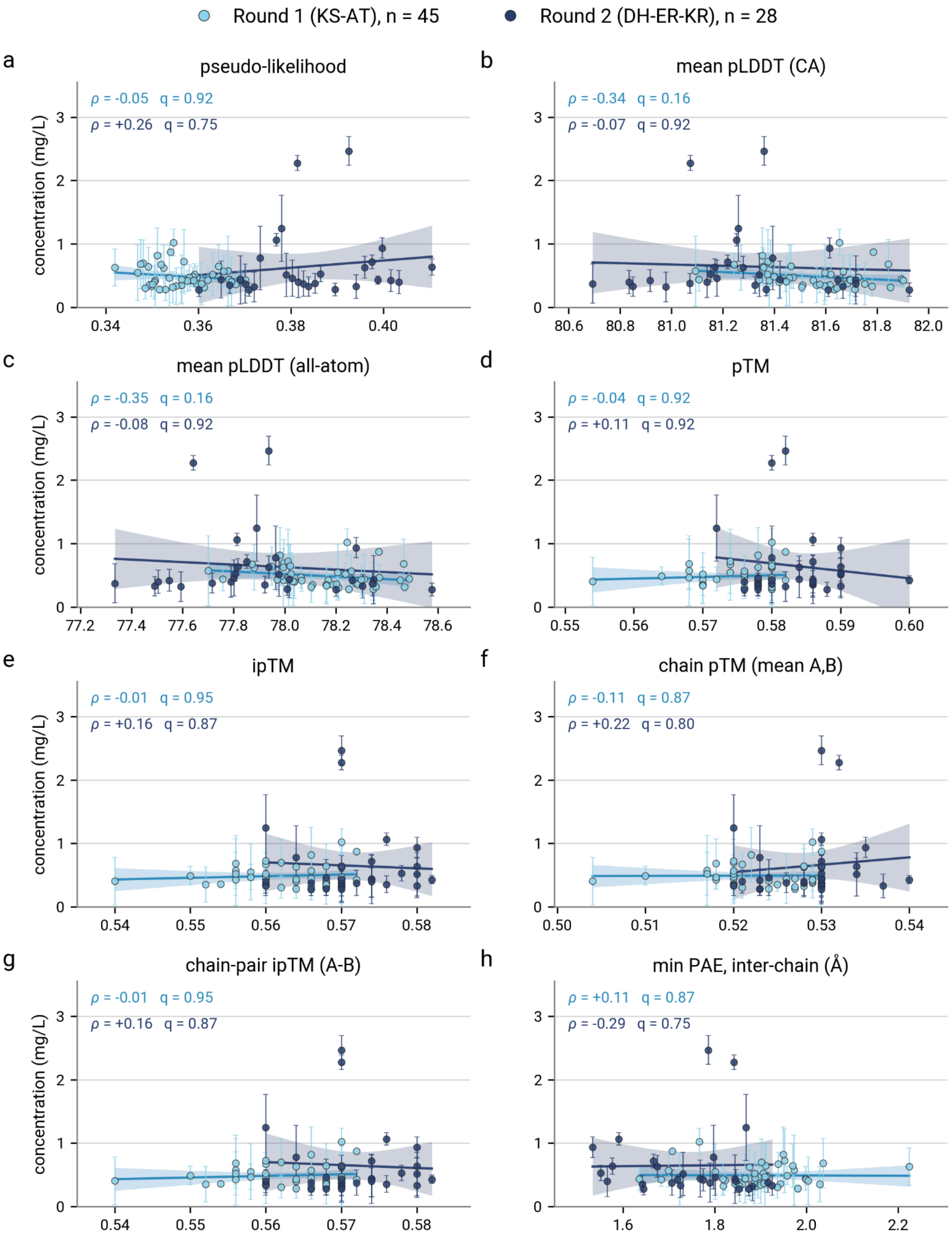

**Figure S15. Relationship between AlphaFold3 confidence and gLM2 Likelihood vs titer.** Each panel plots one score against measured titers for Round 1 (KS-AT, pale blue, n = 45) and Round 2 (DH-ER-KR, navy, n = 28). The line and shaded band are an ordinary-least-squares fit and its 95% mean-prediction interval with Spearman's (ρ) and Benjamini-Hochberg (q) adjusted p-value. **a.** Sequence pseudo-likelihood of each design's own full-module sequence. **b-c.** Mean AlphaFold3 pLDDT over Cα atoms and over all atoms, averaged across five AF3 models and both chains of the homodimer. **d-g.** AlphaFold3's whole-complex and per-chain fit scores: pTM, ipTM, per-chain pTM, and the A–B chain pair ipTM. **h.** Minimum predicted aligned error between the two chains, averaged over both chain orderings.

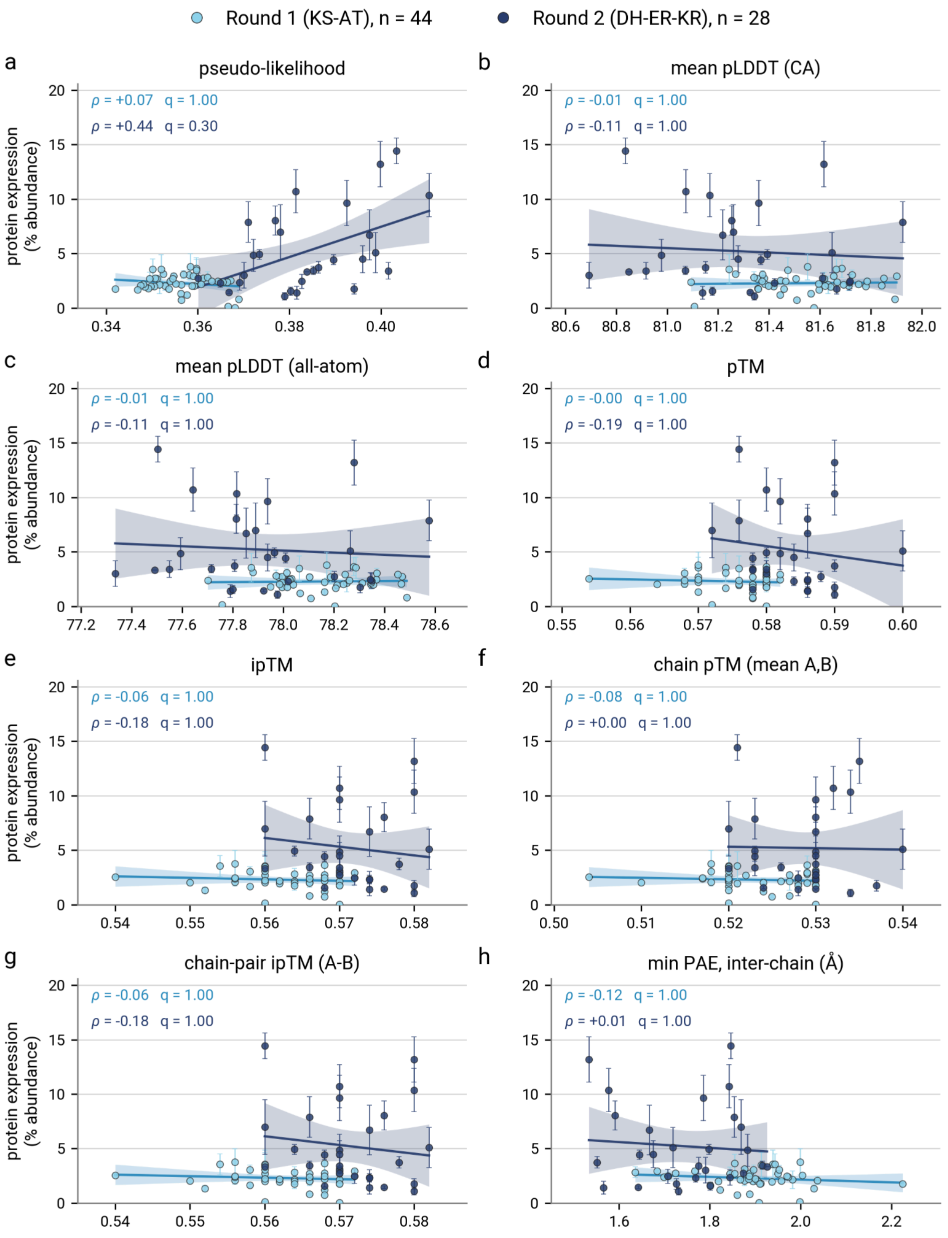

**Figure S16. Relationship between AlphaFold3 confidence and gLM2 Likelihood vs expression.** Each panel plots one score against measured titers for Round 1 (KS-AT, pale blue, n = 45) and Round 2 (DH-ER-KR, navy, n = 28). The line and shaded band are an ordinary-least-squares fit and its 95% mean-prediction interval with Spearman's (ρ) and Benjamini-Hochberg (q) adjusted p-value. **a.** Sequence pseudo-likelihood of each design's own full-module sequence. **b-c.** Mean AlphaFold3 pLDDT over Cα atoms and over all atoms, averaged across five AF3 models and both chains of the homodimer. **d-g.** AlphaFold3's whole-complex and per-chain fit scores: pTM, ipTM, per-chain pTM, and the A–B chain pair ipTM. **h.** Minimum predicted aligned error between the two chains, averaged over both chain orderings.

#### Supplemental Tables 1-16

**Table S1. Boltz-2 PKS co-folding evaluation structures**

| pdb_id | domain | system | method | resolution_A |
|---|---|---|---|---|
| 5XWW | KR | AmphB KR1 G355T/Q364H mutant ternary complex | X-ray | 1.96 |
| 5HV8 | ACP | MLSA2 ACP (mycolactone PKS), octanoyl-phosphopantetheine, solution NMR | solution NMR | |
| 2HFK | TE | Pikromycin TE (PikAIV) + 10-deoxymethynolide product | X-ray | 1.79 |
| 9CGN | TE | Pikromycin TE with covalently trapped linear heptaketide | X-ray | 2.80 |
| 2QO3 | KS | DEBS module 3 KS-AT didomain + cerulenin | X-ray | 2.59 |
| 2H7Y | TE | Pikromycin TE + covalent phosphonate affinity label | X-ray | 2.10 |
| 5D3Z | TE | DEBS TE + allylphosphonate substrate mimic | X-ray | 2.1000 |
| 3MJE | KR | AmphB KR1, NADPH | X-ray | 1.36 |
| 3MJT | KR | AmphB KR1 Q364H mutant, NADPH | X-ray | 1.60 |
| 3MJV | KR | AmphB KR1 W359F mutant, NADPH | X-ray | 1.46 |
| 4L4X | KR | AmphI KR (A2-type), NADPH | X-ray | 2.55 |
| 4HXY | KR | PlmKR1 (phoslactomycin), NADPH | X-ray | 1.68 |
| 5KTK | KR | PksJ KR module 3 (bacillaene), NADPH | X-ray | 1.98 |
| 3SLK | KR-ER | Spinosyn PKS module 2 KR-ER didomain, two NADPH | X-ray | 3.00 |
| 3SLK | KR-ER | Spinosyn PKS module 2 KR-ER didomain, two NADPH | X-ray | 3.00 |
| 8G7W | DH-ER-KR | Type I modPKS reducing region (DH-ER-KR), NADP+ | X-ray | 3.400 |
| 8G7W | DH-ER-KR | Type I modPKS reducing region (DH-ER-KR), NADP+ | X-ray | 3.400 |

**Table S2. Plasmids used in this study.**

| Plasmid | Design | Description | Referenc e | Teselagen ID |
|---|---|---|---|---|
| pJB035_R1-1 | R1-1 | BxbI integration plasmid co-expressing BxbI serine recombinase and R1-1 | This work | fb2b6007-6669-4d2a-84 8c-3b2ac49a1073 |
| pJB035_R1-2 | R1-2 | BxbI integration plasmid co-expressing BxbI serine recombinase and R1-2 | This work | d8db59ff-906f-4d66-8f3 b-e0fd21459431 |
| pJB035_R1-3 | R1-3 | BxbI integration plasmid co-expressing BxbI serine recombinase and R1-3 | This work | fa472b8f-1281-447f-af99 -de4c1c0b7138 |
| pJB035_R1-4 | R1-4 | BxbI integration plasmid co-expressing BxbI serine recombinase and R1-4 | This work | 6ac0a19b-5adf-4bba-a99 c-bc5fd5a57d63 |
| pJB035_R1-6 | R1-6 | BxbI integration plasmid co-expressing BxbI serine recombinase and R1-6 | This work | 373902be-fdec-4d02-aa1 4-7812c22beb6f |
| pJB035_R1-7 | R1-7 | BxbI integration plasmid co-expressing BxbI serine recombinase and R1-7 | This work | 4a63b4e7-da8d-42c9-99 31-65b7f24b6625 |
| pJB035_R1-8 | R1-8 | BxbI integration plasmid co-expressing BxbI serine recombinase and R1-8 | This work | 0f45660b-2af2-42c6-901 9-ccf54b0f4bcf |
| pJB035_R1-9 | R1-9 | BxbI integration plasmid co-expressing BxbI serine recombinase and R1-9 | This work | 2991aabf-78a0-44c5-9e0 5-de48763e0dc5 |
| pJB035_R1-10 | R1-10 | BxbI integration plasmid co-expressing BxbI serine recombinase and R1-10 | This work | f5c056a9-90bc-43b6-983 f-9f2de5e75223 |
| pJB035_R1-11 | R1-11 (R2-Start) | BxbI integration plasmid co-expressing BxbI serine | This work | 12b2f003-4be2-4027-ae1 a-bbb6493a47eb |

| | | | | |
|---|---|---|---|---|
| | | recombinase and R1-11 (R2-Start) | | |
| pJB035_R1-12 | R1-12 | BxbI integration plasmid co-expressing BxbI serine recombinase and R1-12 | This work | 6d356641-4f0a-43f8-81dc-b0e940391d67 |
| pJB035_R1-13 | R1-13 | BxbI integration plasmid co-expressing BxbI serine recombinase and R1-13 | This work | 55f1b8fb-ebc2-49da-9d54-b1ede55346b6 |
| pJB035_R1-14 | R1-14 | BxbI integration plasmid co-expressing BxbI serine recombinase and R1-14 | This work | 4a3f86ff-a65c-453a-8fbb-e23e8516223a |
| pJB035_R1-15 | R1-15 | BxbI integration plasmid co-expressing BxbI serine recombinase and R1-15 | This work | ad26f365-fe14-46ca-b367-d5a043dd0cf8 |
| pJB035_R1-16 | R1-16 | BxbI integration plasmid co-expressing BxbI serine recombinase and R1-16 | This work | d2a143b6-392d-4812-baf2-7ce4b1bbb41d |
| pJB035_R1-17 | R1-17 | BxbI integration plasmid co-expressing BxbI serine recombinase and R1-17 | This work | 1ba0639d-d775-4601-a605-4ce5204e4c4c |
| pJB035_R1-18 | R1-18 | BxbI integration plasmid co-expressing BxbI serine recombinase and R1-18 | This work | 51e6fc88-5fc0-4898-95eb-81c1a9fa8466 |
| pJB035_R1-19 | R1-19 | BxbI integration plasmid co-expressing BxbI serine recombinase and R1-19 | This work | 90a61768-5ec5-45e0-97ef-36ee921a5c42 |
| pJB035_R1-20 | R1-20 | BxbI integration plasmid co-expressing BxbI serine recombinase and R1-20 | This work | a4fe3293-8034-4f39-a761-b73160e8b18f |
| pJB035_R1-21 | R1-21 | BxbI integration plasmid co-expressing BxbI serine recombinase and R1-21 | This work | 74f0366b-ef74-4e47-b708-8d36aa73a5ed |
| pJB035_R1-22 | R1-22 | BxbI integration plasmid co-expressing BxbI serine recombinase and R1-22 | This work | 032ae546-083b-4893-9e7c-c54a02c6a8a7 |

| | | | | |
|---|---|---|---|---|
| pJB035_R1-23 | R1-23 | BxbI integration plasmid co-expressing BxbI serine recombinase and R1-23 | This work | 701de6fe-b66a-47e3-874d-7c40d6a4f7a0 |
| pJB035_R1-24 | R1-24 | BxbI integration plasmid co-expressing BxbI serine recombinase and R1-24 | This work | 305f653d-43de-4883-ac3a-5a265fe2df55 |
| pJB035_R1-25 | R1-25 | BxbI integration plasmid co-expressing BxbI serine recombinase and R1-25 | This work | 50472970-d762-4048-9b03-e9be01e661a6 |
| pJB035_R1-26 | R1-26 | BxbI integration plasmid co-expressing BxbI serine recombinase and R1-26 | This work | aee15c19-4b66-4e24-bdfc-faf0a52b22d2 |
| pJB035_R1-27 | R1-27 | BxbI integration plasmid co-expressing BxbI serine recombinase and R1-27 | This work | 19aeb471-650d-4159-b456-356cd7d911b2 |
| pJB035_R1-28 | R1-28 | BxbI integration plasmid co-expressing BxbI serine recombinase and R1-28 | This work | e3c54754-bea0-4a4a-8e16-135e233d3ef4 |
| pJB035_R1-29 | R1-29 | BxbI integration plasmid co-expressing BxbI serine recombinase and R1-29 | This work | dc3260cb-9de0-4493-9e21-6ac012724cf1 |
| pJB035_R1-30 | R1-30 | BxbI integration plasmid co-expressing BxbI serine recombinase and R1-30 | This work | 26cde64c-4f29-4f4a-8147-4650cbf179b5 |
| pJB035_R1-31 | R1-31 | BxbI integration plasmid co-expressing BxbI serine recombinase and R1-31 | This work | 9aeba43e-1404-42ed-9092-2cf95eac9a95 |
| pJB035_R1-32 | R1-32 | BxbI integration plasmid co-expressing BxbI serine recombinase and R1-32 | This work | c909ab87-50ee-4f18-ad0b-a336dac38a94 |
| pJB035_R1-33 | R1-33 | BxbI integration plasmid co-expressing BxbI serine recombinase and R1-33 | This work | 754cfeb5-9bf6-49f8-8683-cbe5ab3eb724 |

| | | | | |
|---|---|---|---|---|
| pJB035_R1-34 | R1-34 | BxbI integration plasmid co-expressing BxbI serine recombinase and R1-34 | This work | ce7f2c35-cc90-4137-baa8-338df20a612d |
| pJB035_R1-35 | R1-35 | BxbI integration plasmid co-expressing BxbI serine recombinase and R1-35 | This work | 96a27353-1ff0-4e56-b236-9be975cffa87 |
| pJB035_R1-36 | R1-36 | BxbI integration plasmid co-expressing BxbI serine recombinase and R1-36 | This work | 33b46f69-ea3f-45e4-8757-e903754bfee2 |
| pJB035_R1-37 | R1-37 | BxbI integration plasmid co-expressing BxbI serine recombinase and R1-37 | This work | 64ddb302-5268-4438-b085-59899ac3f785 |
| pJB035_R1-40 | R1-40 | BxbI integration plasmid co-expressing BxbI serine recombinase and R1-40 | This work | 5926e890-dac2-4b58-ac92-14e304b2292a |
| pJB035_R1-41 | R1-41 | BxbI integration plasmid co-expressing BxbI serine recombinase and R1-41 | This work | d345d18d-24e8-45dc-ad4f-3a91038178da |
| pJB035_R1-42 | R1-42 | BxbI integration plasmid co-expressing BxbI serine recombinase and R1-42 | This work | 8d861394-0309-4bb6-bf73-c0958c4cadff |
| pJB035_R1-43 | R1-43 | BxbI integration plasmid co-expressing BxbI serine recombinase and R1-43 | This work | 94e865de-e0a8-4083-8214-64dc6192eda1 |
| pJB035_R1-44 | R1-44 | BxbI integration plasmid co-expressing BxbI serine recombinase and R1-44 | This work | 17c6715e-2526-428b-ba07-591203dd4d4b |
| pJB035_R1-45 | R1-45 | BxbI integration plasmid co-expressing BxbI serine recombinase and R1-45 | This work | 299d0550-882f-4e4c-8251-00af520636b6 |
| pJB035_R1-46 | R1-46 | BxbI integration plasmid co-expressing BxbI serine recombinase and R1-46 | This work | 59b3c1ec-0163-49d3-ab2b-a9cf6e3c37ad |

| | | | | |
|---|---|---|---|---|
| pJB035_R1-47 | R1-47 | BxbI integration plasmid co-expressing BxbI serine recombinase and R1-47 | This work | 801851d0-e6b6-4ccf-9691-5924b5773ba0 |
| pJB035_R1-48 | R1-48 | BxbI integration plasmid co-expressing BxbI serine recombinase and R1-48 | This work | be40a91b-f677-429c-b6e2-f246572bd3b2 |
| pJB033_noR4_R2-1 | R2-1 | modified R4 integration plasmid expressing R2-1 | This work | 02ae7263-f70f-49b2-9873-47f56b252c49 |
| pJB033_noR4_R2-2 | R2-2 | modified R4 integration plasmid expressing R2-2 | This work | 24cd6d16-b850-4628-8c3a-110cd302df3e |
| pJB033_noR4_R2-3 | R2-3 | modified R4 integration plasmid expressing R2-3 | This work | 49aad1f0-f2e1-43dd-86be-1450b8f6d7dc |
| pJB033_noR4_R2-4 | R2-4 | modified R4 integration plasmid expressing R2-4 | This work | bfafb9eb-3cee-4d6a-8d90-8b02f4e26e69 |
| pJB033_noR4_R2-5 | R2-5 | modified R4 integration plasmid expressing R2-5 | This work | d2d93e8b-7c26-4037-aedf-e96c4292b1a3 |
| pJB033_noR4_R2-5.1 | R2-5.1 | modified R4 integration plasmid expressing R2-5.1 | This work | 2edfa5c0-490c-467b-897c-b2395e1946fb |
| pJB033_noR4_R2-7 | R2-7 | modified R4 integration plasmid expressing R2-7 | This work | c4907074-cf91-4625-94f3-86e27beeef21 |
| pJB033_noR4_R2-8 | R2-8 | modified R4 integration plasmid expressing R2-8 | This work | 3b616770-2853-4462-95d4-3a2e2d787b92 |
| pJB033_noR4_R2-9 | R2-9 | modified R4 integration plasmid expressing R2-9 | This work | af2caf4b-1783-4639-9d14-61fda3c98b6d |
| pJB033_noR4_R2-11 | R2-11 | modified R4 integration plasmid expressing R2-11 | This work | 9a0de9bf-96fd-4b90-ac9e-a3d824b88c1f |
| pJB033_noR4_R2-11.1 | R2-11.1 | modified R4 integration plasmid expressing R2-11.1 | This work | ea8386cc-5848-4887-8989-9cc49bb2c982 |
| pJB033_noR4_R2-12 | R2-12 | modified R4 integration plasmid expressing R2-12 | This work | 0b423a49-c68b-4045-bf72-0d3c5ad603b0 |
| pJB033_noR4_R2-12.1 | R2-12.1 | modified R4 integration plasmid expressing R2-12.1 | This work | c92dd359-d519-47d3-a879-cd7463645333 |

| | | | | |
|---|---|---|---|---|
| pJB033_noR4_ R2-14 | R2-14 | modified R4 integration plasmid expressing R2-14 | This work | fa712792-6d95-4972-96 6f-9ad3bf1777d7 |
| pJB033_noR4_ R2-15 | R2-15 | modified R4 integration plasmid expressing R2-15 | This work | 5e387464-86a0-4646-b2 b7-3048b2894b10 |
| pJB033_noR4_ R2-15.1 | R2-15.1 | modified R4 integration plasmid expressing R2-15.1 | This work | f632a772-2377-4832-83e d-d53a75472db5 |
| pJB033_noR4_ R2-17 | R2-17 | modified R4 integration plasmid expressing R2-17 | This work | fc04492e-7d6c-48d9-89e 9-a4b69680704d |
| pJB033_noR4_ R2-17.1 | R2-17.1 | modified R4 integration plasmid expressing R2-17.1 | This work | 815d1244-0bfa-4718-94 68-7604755978a6 |
| pJB033_noR4_ R2-17.2 | R2-17.2 | modified R4 integration plasmid expressing R2-17.2 | This work | f3a970e1-d560-4fcb-a8a 3-78abf80c1b78 |
| pJB033_noR4_ R2-17.3 | R2-17.3 | modified R4 integration plasmid expressing R2-17.3 | This work | 4f967644-285b-4f1c-8b2 7-d51f28be1dfc |
| pJB033_noR4_ R2-21 | R2-21 | modified R4 integration plasmid expressing R2-21 | This work | 4f80205d-f18f-43a7-ae0 0-2994adbde3a4 |
| pJB033_noR4_ R2-22 | R2-22 | modified R4 integration plasmid expressing R2-22 | This work | fe5c8f89-4d2d-44dc-9eb 0-04fc200b12d1 |
| pJB033_noR4_ R2-23 | R2-23 | modified R4 integration plasmid expressing R2-23 | This work | 6641d8a8-6a01-46e6-98 cf-aeb7d55de233 |
| pJB033_noR4_ R2-23.1 | R2-23.1 | modified R4 integration plasmid expressing R2-23.1 | This work | 5d34b014-cf0e-4303-bb3 b-e45a2befca14 |
| pJB033_noR4_ R2-23.2 | R2-23.2 | modified R4 integration plasmid expressing R2-23.2 | This work | 6ced0780-4fb5-4e2a-aef 3-92cd8eb87f19 |
| pJB033_noR4_ R2-26 | R2-26 | modified R4 integration plasmid expressing R2-26 | This work | c30c8dd8-a6ae-4a9c-812 e-1cf970a60ddc |
| pJB033_noR4_ R2-26.1 | R2-26.1 | modified R4 integration plasmid expressing R2-26.1 | This work | 931bd0b5-2cef-4762-a61 f-01ea98e1c08a |
| pJB033_noR4_ R2-26.2 | R2-26.2 | modified R4 integration plasmid expressing R2-26.2 | This work | 329ceff3-5b51-4aa5-aef0 -0102193c4b70 |
| pJB033_noR4_ R2-29 | R2-29 | modified R4 integration plasmid expressing R2-29 | This work | 3079a3b6-8840-4719-b6 11-15609d57bff1 |

| | | | | |
|---|---|---|---|---|
| pJB033_noR4_ R2-30 | R2-30 | modified R4 integration plasmid expressing R2-30 | This work | 759bcc38-3f58-418b-b374-91724392e9b2 |
| pJB033_noR4_ R2-31 | R2-31 | modified R4 integration plasmid expressing R2-31 | This work | 8fb668dc-a714-436e-bf5c-ff8b259c1384 |
| pJB033_noR4_ R2-32 | R2-32 | modified R4 integration plasmid expressing R2-32 | This work | 8b840f98-f348-47c0-a19c-c9fb06282967 |
| pJB033_noR4_ R2-33 | R2-33 | modified R4 integration plasmid expressing R2-33 | This work | af862e37-9937-48a2-ab5e-e5d8b7406bb8 |
| pJB033_noR4_ R2-34 | R2-34 | modified R4 integration plasmid expressing R2-34 | This work | 7fb5d543-db8f-4cfd-ad29-e6d7153d9e9c |
| pJB033_noR4_ R2-35 | R2-35 | modified R4 integration plasmid expressing R2-35 | This work | 5840032f-6653-4733-81c7-97d5424da960 |
| pJB033_noR4_ R2-36 | R2-36 | modified R4 integration plasmid expressing R2-36 | This work | e1474976-0e4a-4f0a-b3a5-d396ab8273a1 |
| pJB033_noR4_ R2-36.1 | R2-36.1 | modified R4 integration plasmid expressing R2-36.1 | This work | 49958000-d868-4cf3-92d4-bdf770250996 |
| pJB033_noR4_ R2-38 | R2-38 | modified R4 integration plasmid expressing R2-38 | This work | 4852a1c2-2836-470c-8b76-55859b2c8388 |
| pJB033_noR4_ R2-39 | R2-39 | modified R4 integration plasmid expressing R2-39 | This work | 6bbf969f-867d-4078-8dc8-9c9156f86015 |
| pJB033_noR4_ R2-40 | R2-40 | modified R4 integration plasmid expressing R2-40 | This work | a25431f3-398a-4dc1-8c70-d76d84031efe |
| pJB033_noR4_ R2-40.1 | R2-40.1 | modified R4 integration plasmid expressing R2-40.1 | This work | 2519dbb5-08e1-4afb-8147-133a573e7730 |
| pJB033_noR4_ R1-11 | R1-11 (R2-Start) | modified R4 integration plasmid expressing R1-11 (R2-Start) | This work | 96082da1-b1c3-4810-8b12-a7d057b6039d |
| pJB033_noR4_ R2-starter | R2 starter | modified R4 integration plasmid encoding R2-Start with inserted BsmBI recognition sites for Golden Gate assembly | This work | 4dbfef9e-c3b9-4a9f-895c-2af5fbc2b8ea |

| | | | | |
|---|---|---|---|---|
| pJB035_R1-starter | R1 starter | BxbI integration plasmid co-expressing BxbI serine recombinase and encoding VL-Start with inserted BsmBI recognition sites for Golden Gate assembly | This work | d9373b1e-bb1e-4043-9ea0-f838b01af34e |

**Table S3. Strains used in this study.**

| Species | Strain Name | Design | PKS integration locus | Description | Reference | Teselagen Strain ID |
|---|---|---|---|---|---|---|
| *E. coli* | NEB Stables | | | NEB | | |
| *P. putida* | LM3 | | | RV::pAN002 LM3 | Lee *et al*[24] | 7f08450c-50f4-4ef3-8b61-e3ff11aa413d |
| *P. putida* | LM3-FlvJ R1-1 | R1-1 | BxB1 | RV::pAN002 LM3 TG1::FlvJ BxB1::pJB035_R1-1 | This work | b3aa5fdf-e293-466a-aa6b-bd4e9cd36db9 |
| *P. putida* | LM3-FlvJ R1-2 | R1-2 | BxB1 | RV::pAN002 LM3 TG1::FlvJ BxB1::pJB035_R1-2 | This work | c6f7def5-5c7c-4493-8a18-40a40b06130b |
| *P. putida* | LM3-FlvJ R1-3 | R1-3 | BxB1 | RV::pAN002 LM3 TG1::FlvJ BxB1::pJB035_R1-3 | This work | 063c5dd9-9e1c-4ed1-b5b5-4745c9a032fe |
| *P. putida* | LM3-FlvJ R1-4 | R1-4 | BxB1 | RV::pAN002 LM3 TG1::FlvJ BxB1::pJB035_R1-4 | This work | 4bdc8abf-af96-459b-947b-94af718775ea |
| *P. putida* | LM3-FlvJ R1-6 | R1-6 | BxB1 | RV::pAN002 LM3 TG1::FlvJ BxB1::pJB035_R1-6 | This work | f797e3db-a2a1-43f6-8013- |

| | | | | | | |
|---|---|---|---|---|---|---|
| | | | | | | d9b029e15c0d |
| *P. putida* | LM3-FlvJ R1-7 | R1-7 | BxB1 | RV::pAN002 LM3 TG1::FlvJ BxB1::pJB035_R1-7 | This work | 3147678d-0791-43dd-9864-263021b98c7b |
| *P. putida* | LM3-FlvJ R1-8 | R1-8 | BxB1 | RV::pAN002 LM3 TG1::FlvJ BxB1::pJB035_R1-8 | This work | 377a8dfa-067c-4a6b-97c2-2eb9b936a452 |
| *P. putida* | LM3-FlvJ R1-9 | R1-9 | BxB1 | RV::pAN002 LM3 TG1::FlvJ BxB1::pJB035_R1-9 | This work | 69e557c3-93c4-49e0-b435-9335589f635a |
| *P. putida* | LM3-FlvJ R1-10 | R1-10 | BxB1 | RV::pAN002 LM3 TG1::FlvJ BxB1::pJB035_R1-10 | This work | 05f14858-fdc2-4e73-af3b-080f96314c6a |
| *P. putida* | LM3-FlvJ R1-11 (R2-Start) | R1-11 (R2-Start) | BxB1 | RV::pAN002 LM3 TG1::FlvJ BxB1::pJB035_R1-11 | This work | 38a3850a-461c-4819-b3d7-7da724effaf8 |
| *P. putida* | LM3-FlvJ R1-12 | R1-12 | BxB1 | RV::pAN002 LM3 TG1::FlvJ BxB1::pJB035_R1-12 | This work | 64f8cb12-76a9-480d-bb10-ce34768cb32c |
| *P. putida* | LM3-FlvJ R1-13 | R1-13 | BxB1 | RV::pAN002 LM3 TG1::FlvJ BxB1::pJB035_R1-13 | This work | 0512e7c9-b72b-4e72-a05c-e8128551379f |
| *P. putida* | LM3-FlvJ R1-14 | R1-14 | BxB1 | RV::pAN002 LM3 TG1::FlvJ BxB1::pJB035_R1-14 | This work | 5573272d-9c0c-4a7d-93a7-28a8b5c15019 |

| | | | | | | |
|---|---|---|---|---|---|---|
| *P. putida* | LM3-FlvJ R1-15 | R1-15 | BxB1 | RV::pAN002 LM3 TG1::FlvJ BxB1::pJB035_R1-15 | This work | e741f7d3-dc8b-4697-90e1-cfc91d0b0948 |
| *P. putida* | LM3-FlvJ R1-16 | R1-16 | BxB1 | RV::pAN002 LM3 TG1::FlvJ BxB1::pJB035_R1-16 | This work | e407dceb-bd64-4b24-8f0c-f4be6b8868a2 |
| *P. putida* | LM3-FlvJ R1-17 | R1-17 | BxB1 | RV::pAN002 LM3 TG1::FlvJ BxB1::pJB035_R1-17 | This work | 2852772e-f93d-4bcd-8cea-76acf1d71612 |
| *P. putida* | LM3-FlvJ R1-18 | R1-18 | BxB1 | RV::pAN002 LM3 TG1::FlvJ BxB1::pJB035_R1-18 | This work | dd5039cc-8323-4277-b5b1-ac08616d415c |
| *P. putida* | LM3-FlvJ R1-19 | R1-19 | BxB1 | RV::pAN002 LM3 TG1::FlvJ BxB1::pJB035_R1-19 | This work | 1499338f-486e-40f1-b354-97569d26c945 |
| *P. putida* | LM3-FlvJ R1-20 | R1-20 | BxB1 | RV::pAN002 LM3 TG1::FlvJ BxB1::pJB035_R1-20 | This work | 2f0f341a-26ad-492d-b086-eb74c8ffa7fe |
| *P. putida* | LM3-FlvJ R1-21 | R1-21 | BxB1 | RV::pAN002 LM3 TG1::FlvJ BxB1::pJB035_R1-21 | This work | a7b339ac-f4d7-4195-87eb-638524b5ac0e |
| *P. putida* | LM3-FlvJ R1-22 | R1-22 | BxB1 | RV::pAN002 LM3 TG1::FlvJ BxB1::pJB035_R1-22 | This work | 459902a8-cf2c-43d8-ab4c-e9e80ba32599 |
| *P. putida* | LM3-FlvJ R1-23 | R1-23 | BxB1 | RV::pAN002 LM3 TG1::FlvJ BxB1::pJB035_R1-23 | This work | db650756-7903-4dfa-ab5c-b9b92ce93a30 |

| | | | | | | |
|---|---|---|---|---|---|---|
| *P. putida* | LM3-FlvJ R1-24 | R1-24 | BxB1 | RV::pAN002 LM3 TG1::FlvJ BxB1::pJB035_R1-24 | This work | 04089647-2036-448b-915e-3763650fc9fa |
| *P. putida* | LM3-FlvJ R1-25 | R1-25 | BxB1 | RV::pAN002 LM3 TG1::FlvJ BxB1::pJB035_R1-25 | This work | 12b47cc6-9faf-427f-97e4-f2abf1f71b8e |
| *P. putida* | LM3-FlvJ R1-26 | R1-26 | BxB1 | RV::pAN002 LM3 TG1::FlvJ BxB1::pJB035_R1-26 | This work | 25076e65-617e-4bb6-9c0a-fe5bb5f9282a |
| *P. putida* | LM3-FlvJ R1-27 | R1-27 | BxB1 | RV::pAN002 LM3 TG1::FlvJ BxB1::pJB035_R1-27 | This work | 6e77a671-9e5f-41e6-87e6-c72caf1e1dfe |
| *P. putida* | LM3-FlvJ R1-28 | R1-28 | BxB1 | RV::pAN002 LM3 TG1::FlvJ BxB1::pJB035_R1-28 | This work | 87244984-eaa6-42b9-8f6a-c4e9b559bdc2 |
| *P. putida* | LM3-FlvJ R1-29 | R1-29 | BxB1 | RV::pAN002 LM3 TG1::FlvJ BxB1::pJB035_R1-29 | This work | 5e5d4bd0-0767-484a-ba3a-a2076c7a2f83 |
| *P. putida* | LM3-FlvJ R1-30 | R1-30 | BxB1 | RV::pAN002 LM3 TG1::FlvJ BxB1::pJB035_R1-30 | This work | 9cd50094-174d-432b-8cbb-6d12c35f7f79 |
| *P. putida* | LM3-FlvJ R1-31 | R1-31 | BxB1 | RV::pAN002 LM3 TG1::FlvJ BxB1::pJB035_R1-31 | This work | 73379f98-1279-4c95-b943-e1745fee3e3c |
| *P. putida* | LM3-FlvJ R1-32 | R1-32 | BxB1 | RV::pAN002 LM3 TG1::FlvJ BxB1::pJB035_R1-32 | This work | cef910f5-6ffe-4db1-8244-be5e27f7fa4f |

| | | | | | | |
|---|---|---|---|---|---|---|
| *P. putida* | LM3-FlvJ R1-33 | R1-33 | BxB1 | RV::pAN002 LM3 TG1::FlvJ BxB1::pJB035_R1-33 | This work | 262d50b2-aca3-4e79-ac13-8c76586569ba |
| *P. putida* | LM3-FlvJ R1-34 | R1-34 | BxB1 | RV::pAN002 LM3 TG1::FlvJ BxB1::pJB035_R1-34 | This work | 693e12f1-676b-45fb-8650-79ac0e4f8861 |
| *P. putida* | LM3-FlvJ R1-35 | R1-35 | BxB1 | RV::pAN002 LM3 TG1::FlvJ BxB1::pJB035_R1-35 | This work | b92f45fe-e040-4543-bf82-7f25162cd376 |
| *P. putida* | LM3-FlvJ R1-36 | R1-36 | BxB1 | RV::pAN002 LM3 TG1::FlvJ BxB1::pJB035_R1-36 | This work | 92add436-6e6f-4efa-896c-0dedce572d7e |
| *P. putida* | LM3-FlvJ R1-37 | R1-37 | BxB1 | RV::pAN002 LM3 TG1::FlvJ BxB1::pJB035_R1-37 | This work | 93574e5f-99cb-4a46-b1b8-a5ba9662d1d4 |
| *P. putida* | LM3-FlvJ R1-40 | R1-40 | BxB1 | RV::pAN002 LM3 TG1::FlvJ BxB1::pJB035_R1-40 | This work | b8c25437-8c70-426f-8cf1-d77ddf4f5e3e |
| *P. putida* | LM3-FlvJ R1-41 | R1-41 | BxB1 | RV::pAN002 LM3 TG1::FlvJ BxB1::pJB035_R1-41 | This work | b17ab48a-7fb9-4b65-a4a7-b165bdfff24a |
| *P. putida* | LM3-FlvJ R1-42 | R1-42 | BxB1 | RV::pAN002 LM3 TG1::FlvJ BxB1::pJB035_R1-42 | This work | 9ced70ca-fa15-4f32-b9e9-ca7fb18fca00 |
| *P. putida* | LM3-FlvJ R1-43 | R1-43 | BxB1 | RV::pAN002 LM3 TG1::FlvJ BxB1::pJB035_R1-43 | This work | 00f7ea94-34d6-4c09-be40-84af2dfb1bf9 |

| | | | | | | |
|---|---|---|---|---|---|---|
| *P. putida* | LM3-FlvJ R1-44 | R1-44 | BxB1 | RV::pAN002 LM3 TG1::FlvJ BxB1::pJB035_R1-44 | This work | 308b010c-c5db-464a-8706-c97273a30688 |
| *P. putida* | LM3-FlvJ R1-45 | R1-45 | BxB1 | RV::pAN002 LM3 TG1::FlvJ BxB1::pJB035_R1-45 | This work | 44e9a61c-98b9-4705-bb22-6f8935de7620 |
| *P. putida* | LM3-FlvJ R1-46 | R1-46 | BxB1 | RV::pAN002 LM3 TG1::FlvJ BxB1::pJB035_R1-46 | This work | 4dc1eba7-b27a-4ebb-b8b5-02942126e4d2 |
| *P. putida* | LM3-FlvJ R1-47 | R1-47 | BxB1 | RV::pAN002 LM3 TG1::FlvJ BxB1::pJB035_R1-47 | This work | 9b34569b-1e04-4a40-a16a-0d3544e044a5 |
| *P. putida* | LM3-FlvJ R1-48 | R1-48 | BxB1 | RV::pAN002 LM3 TG1::FlvJ BxB1::pJB035_R1-48 | This work | e5b13e87-4232-4ed7-adfa-227ce377dd37 |
| *P. putida* | LM3-FlvJ R2-1 | R2-1 | R4 | RV::pAN002 LM3 TG1::FlvJ R4::pJB033_noR4_R2-1 | This work | fa637e31-2be0-42b1-b295-bc75ea359479 |
| *P. putida* | LM3-FlvJ R2-2 | R2-2 | R4 | RV::pAN002 LM3 TG1::FlvJ R4::pJB033_noR4_R2-2 | This work | 67832c17-d9c6-4b3c-af12-5a104027277f |
| *P. putida* | LM3-FlvJ R2-3 | R2-3 | R4 | RV::pAN002 LM3 TG1::FlvJ R4::pJB033_noR4_R2-3 | This work | 2d69ec81-cdb4-4244-983e-3c7eb9744250 |
| *P. putida* | LM3-FlvJ R2-4 | R2-4 | R4 | RV::pAN002 LM3 TG1::FlvJ R4::pJB033_noR4_R2-4 | This work | 4b4077fb-c0ee-4d0d-96f7-06db1c1baeb7 |

| | | | | | | |
|---|---|---|---|---|---|---|
| *P. putida* | LM3-FlvJ R2-5 | R2-5 | R4 | RV::pAN002 LM3 TG1::FlvJ R4::pJB033_noR4_R2-5 | This work | e939ad9d-8394-4e39-a9d5-97a2ea097921 |
| *P. putida* | LM3-FlvJ R2-5.1 | R2-5.1 | R4 | RV::pAN002 LM3 TG1::FlvJ R4::pJB033_noR4_R2-5.1 | This work | 8f24cc15-e3e4-46fa-ac39-f3832362f237 |
| *P. putida* | LM3-FlvJ R2-7 | R2-7 | R4 | RV::pAN002 LM3 TG1::FlvJ R4::pJB033_noR4_R2-7 | This work | b0ff6412-b4c5-4741-8e58-5da861a7fc2f |
| *P. putida* | LM3-FlvJ R2-8 | R2-8 | R4 | RV::pAN002 LM3 TG1::FlvJ R4::pJB033_noR4_R2-8 | This work | 076626e0-5324-4f06-876a-98176e714997 |
| *P. putida* | LM3-FlvJ R2-9 | R2-9 | R4 | RV::pAN002 LM3 TG1::FlvJ R4::pJB033_noR4_R2-9 | This work | ba70c5cb-c3bb-4a05-9ab5-2883fa26da0f |
| *P. putida* | LM3-FlvJ R2-11 | R2-11 | R4 | RV::pAN002 LM3 TG1::FlvJ R4::pJB033_noR4_R2-11 | This work | 09475f84-b23a-4368-b60d-989265bf0471 |
| *P. putida* | LM3-FlvJ R2-11.1 | R2-11.1 | R4 | RV::pAN002 LM3 TG1::FlvJ R4::pJB033_noR4_R2-11.1 | This work | e7229829-be97-4587-84f7-4d4cc3d4f360 |
| *P. putida* | LM3-FlvJ R2-12 | R2-12 | R4 | RV::pAN002 LM3 TG1::FlvJ R4::pJB033_noR4_R2-12 | This work | 7db8c837-1d68-4cfa-9994-50afb3f7b845 |
| *P. putida* | LM3-FlvJ R2-12.1 | R2-12.1 | R4 | RV::pAN002 LM3 TG1::FlvJ R4::pJB033_noR4_R2-12.1 | This work | 73c1145f-1261-4924-894b-1995cc2ccd4e |

| | | | | | | |
|---|---|---|---|---|---|---|
| *P. putida* | LM3-FlvJ R2-14 | R2-14 | R4 | RV::pAN002 LM3 TG1::FlvJ R4::pJB033_noR4_R2-14 | This work | 406083e4-8a22-4ee0-a4b0-2702b935b218 |
| *P. putida* | LM3-FlvJ R2-15 | R2-15 | R4 | RV::pAN002 LM3 TG1::FlvJ R4::pJB033_noR4_R2-15 | This work | b691ec8c-bbd6-4814-9615-dc844474e64a |
| *P. putida* | LM3-FlvJ R2-15.1 | R2-15.1 | R4 | RV::pAN002 LM3 TG1::FlvJ R4::pJB033_noR4_R2-15.1 | This work | b98e4f27-de64-4bee-9f56-5b195a8b0c18 |
| *P. putida* | LM3-FlvJ R2-17 | R2-17 | R4 | RV::pAN002 LM3 TG1::FlvJ R4::pJB033_noR4_R2-17 | This work | 423868f8-f002-47f1-836a-040d7ac92e7f |
| *P. putida* | LM3-FlvJ R2-17.1 | R2-17.1 | R4 | RV::pAN002 LM3 TG1::FlvJ R4::pJB033_noR4_R2-17.1 | This work | 0964a031-4e60-4410-83ca-3cf4593471a6 |
| *P. putida* | LM3-FlvJ R2-17.2 | R2-17.2 | R4 | RV::pAN002 LM3 TG1::FlvJ R4::pJB033_noR4_R2-17.2 | This work | 9f2e4e56-2711-4461-999c-fe4995f83238 |
| *P. putida* | LM3-FlvJ R2-17.3 | R2-17.3 | R4 | RV::pAN002 LM3 TG1::FlvJ R4::pJB033_noR4_R2-17.3 | This work | 1c94407f-59d8-4ebc-bae6-f6ef589d01bd |
| *P. putida* | LM3-FlvJ R2-21 | R2-21 | R4 | RV::pAN002 LM3 TG1::FlvJ R4::pJB033_noR4_R2-21 | This work | 7078b182-b2ba-4a36-8a55-93d54654844d |
| *P. putida* | LM3-FlvJ R2-22 | R2-22 | R4 | RV::pAN002 LM3 TG1::FlvJ R4::pJB033_noR4_R2-22 | This work | 9d896378-2158-4b33-90c8-30461eac3bb7 |

| | | | | | | |
|---|---|---|---|---|---|---|
| *P. putida* | LM3-FlvJ R2-23 | R2-23 | R4 | RV::pAN002 LM3 TG1::FlvJ R4::pJB033_noR4_R2-23 | This work | 2361e0f5-f8e3-4745-82dc-8e4421f1af2b |
| *P. putida* | LM3-FlvJ R2-23.1 | R2-23.1 | R4 | RV::pAN002 LM3 TG1::FlvJ R4::pJB033_noR4_R2-23.1 | This work | 6a275a91-3a60-4a86-bcd0-9d9d9c4c9160 |
| *P. putida* | LM3-FlvJ R2-23.2 | R2-23.2 | R4 | RV::pAN002 LM3 TG1::FlvJ R4::pJB033_noR4_R2-23.2 | This work | 5c80bd2b-821a-4c9e-8dc0-4029be491815 |
| *P. putida* | LM3-FlvJ R2-26 | R2-26 | R4 | RV::pAN002 LM3 TG1::FlvJ R4::pJB033_noR4_R2-26 | This work | 95549bcf-bf21-45ef-96f3-40836c8b5bca |
| *P. putida* | LM3-FlvJ R2-26.1 | R2-26.1 | R4 | RV::pAN002 LM3 TG1::FlvJ R4::pJB033_noR4_R2-26.1 | This work | 740356cf-bd66-40d6-9641-12e1b4da27bc |
| *P. putida* | LM3-FlvJ R2-26.2 | R2-26.2 | R4 | RV::pAN002 LM3 TG1::FlvJ R4::pJB033_noR4_R2-26.2 | This work | a6f2a1fa-6d5c-4858-bbdc-3abf184a5241 |
| *P. putida* | LM3-FlvJ R2-29 | R2-29 | R4 | RV::pAN002 LM3 TG1::FlvJ R4::pJB033_noR4_R2-29 | This work | fb17ffab-b303-4067-a438-39fa602a15b9 |
| *P. putida* | LM3-FlvJ R2-30 | R2-30 | R4 | RV::pAN002 LM3 TG1::FlvJ R4::pJB033_noR4_R2-30 | This work | 2c25d5d7-e0d8-40e0-9a61-6845d723c097 |
| *P. putida* | LM3-FlvJ R2-31 | R2-31 | R4 | RV::pAN002 LM3 TG1::FlvJ R4::pJB033_noR4_R2-31 | This work | 59aaebd8-7268-4b75-9627-5a44ee8c5a7c |

| | | | | | | |
|---|---|---|---|---|---|---|
| *P. putida* | LM3-FlvJ R2-32 | R2-32 | R4 | RV::pAN002 LM3 TG1::FlvJ R4::pJB033_noR4_R2-32 | This work | f5849a28-efb1-4be6-8acd-88e86f868b56 |
| *P. putida* | LM3-FlvJ R2-33 | R2-33 | R4 | RV::pAN002 LM3 TG1::FlvJ R4::pJB033_noR4_R2-33 | This work | 9a267314-3138-4378-b7b4-f384a3c4497b |
| *P. putida* | LM3-FlvJ R2-34 | R2-34 | R4 | RV::pAN002 LM3 TG1::FlvJ R4::pJB033_noR4_R2-34 | This work | accb12da-16b1-4787-88fe-b597a94dbbc8 |
| *P. putida* | LM3-FlvJ R2-35 | R2-35 | R4 | RV::pAN002 LM3 TG1::FlvJ R4::pJB033_noR4_R2-35 | This work | 2c769c8a-62c5-48e2-ba0d-05585b46bc44 |
| *P. putida* | LM3-FlvJ R2-36 | R2-36 | R4 | RV::pAN002 LM3 TG1::FlvJ R4::pJB033_noR4_R2-36 | This work | 4367e9f5-417c-4457-866f-bceb683e2bfc |
| *P. putida* | LM3-FlvJ R2-36.1 | R2-36.1 | R4 | RV::pAN002 LM3 TG1::FlvJ R4::pJB033_noR4_R2-36.1 | This work | dd972a5c-ed3e-4a50-9618-4f65f94003c4 |
| *P. putida* | LM3-FlvJ R2-38 | R2-38 | R4 | RV::pAN002 LM3 TG1::FlvJ R4::pJB033_noR4_R2-38 | This work | 23a9228e-d66b-4eb7-8d82-d1d4580b38bd |
| *P. putida* | LM3-FlvJ R2-39 | R2-39 | R4 | RV::pAN002 LM3 TG1::FlvJ R4::pJB033_noR4_R2-39 | This work | b379e969-ebdb-410e-9022-3d32cba6561e |
| *P. putida* | LM3-FlvJ R2-40 | R2-40 | R4 | RV::pAN002 LM3 TG1::FlvJ R4::pJB033_noR4_R2-40 | This work | 5f3b28e9-f4db-4835-a9b3-d61b67c5be98 |

| | | | | | | |
|---|---|---|---|---|---|---|
| *P. putida* | LM3-FlvJ R2-40.1 | R2-40.1 | R4 | RV::pAN002 LM3 TG1::FlvJ R4::pJB033_noR4_R2-40.1 | This work | 36de216b-fe52-4011-9309-def1d4bf2378 |
| *P. putida* | LM3-FlvJ cmipks163 (Lee et al strain) | Lee et al | BxB1 | RV::pAN002 LM3 TG1::FlvJ BxB1::pBH026_CmiPksM2433L | Lee *et al*[24] | 7a04c3dd-cacc-47b0-b668-d48add4f7e3c |
| *P. putida* | LM3-FlvJ | base strain | | RV::pAN002 LM3 TG1::FlvJ | Lee *et al*[24] | 43e32dbb-6c63-4ec8-ba54-c7cf5f978800 |
| *P. putida* | LM3-FlvJ R1-11 (R2-Start) | R1-11 (R2-Start) | R4 | RV::pAN002 LM3 TG1::FlvJ R4::pJB033_noR4_R1-11 | This work | 490f3675-e1ac-430a-81f5-6719e1b69083 |
| *P. putida* | LM3-FlvJ cmipks163 (VL-Start) | VL-Start | BxB1 | RV::pAN002 LM3 TG1::FlvJ BxB1::pjb035_cmipks163 | This work | 1b71492a-4ef3-41af-b920-2c54b389fdfb |

#### Supplementary References

1. Klass, S. H. *et al.* Engineering an extremely hybrid PKS for adipic acid production. *ACS Synth. Biol.* **15**, 2887–2899 (2026).
2. Leesong, M., Henderson, B. S., Gillig, J. R., Schwab, J. M. & Smith, J. L. Structure of a dehydratase-isomerase from the bacterial pathway for biosynthesis of unsaturated fatty acids: two catalytic activities in one active site. *Structure* **4**, 253–264 (1996).
3. Akey, D. L. *et al.* Crystal structures of dehydratase domains from the curacin polyketide biosynthetic pathway. *Structure* **18**, 94–105 (2010).
4. Tsai, S.-C. S. & Ames, B. D. Structural enzymology of polyketide synthases. *Methods Enzymol.* **459**, 17–47 (2009).
5. Keatinge-Clay, A. T. A tylosin ketoreductase reveals how chirality is determined in polyketides. *Chem. Biol.* **14**, 898–908 (2007).
6. Keatinge-Clay, A. T. & Stroud, R. M. The structure of a ketoreductase determines the organization of the beta-carbon processing enzymes of modular polyketide synthases. *Structure* **14**, 737–748 (2006).
7. Maier, T., Leibundgut, M. & Ban, N. The crystal structure of a mammalian fatty acid synthase. *Science* **321**, 1315–1322 (2008).
8. Zheng, J., Gay, D. C., Demeler, B., White, M. A. & Keatinge-Clay, A. T. Divergence of multimodular polyketide synthases revealed by a didomain structure. *Nat. Chem. Biol.* **8**, 615–621 (2012).
9. Kwan, D. H. & Leadlay, P. F. Mutagenesis of a modular polyketide synthase enoylreductase domain reveals insights into catalysis and stereospecificity. *ACS Chem. Biol.* **5**, 829–838 (2010).
10. Kwan, D. H. *et al.* Prediction and manipulation of the stereochemistry of enoylreduction in modular polyketide synthases. *Chem. Biol.* **15**, 1231–1240 (2008).

11. Tang, Y., Kim, C.-Y., Mathews, I. I., Cane, D. E. & Khosla, C. The 2.7-Å crystal structure of a 194-kDa homodimeric fragment of the 6-deoxyerythronolide B synthase. *Proc. Natl. Acad. Sci. U. S. A.* **103**, 11124–11129 (2006).

12. McCullough, T. M. *et al.* Structure of a modular polyketide synthase reducing region. *Structure* **31**, 1109-1120.e3 (2023).

13. Bagde, S. R., Mathews, I. I., Fromme, J. C. & Kim, C.-Y. Modular polyketide synthase contains two reaction chambers that operate asynchronously. *Science* **374**, 723–729 (2021).

14. Yuzawa, S. *et al.* Comprehensive in vitro analysis of acyltransferase domain exchanges in modular polyketide synthases and its application for short-chain ketone production. *ACS Synth. Biol.* **6**, 139–147 (2017).

15. Lee, N. *et al.* Retrobiosynthesis of unnatural lactams via reprogrammed polyketide synthase. *Nat. Catal.* **8**, 389–402 (2025).
